# ALC1 links chromatin accessibility to PARP inhibitor response in homologous recombination deficient cells

**DOI:** 10.1101/2020.12.16.422851

**Authors:** Priyanka Verma, Yeqiao Zhou, Zhendong Cao, Peter V. Deraska, Moniher Deb, Eri Arai, Weihua Li, Yue Shao, Yiwen Li, Laura Puentes, Sonali Patankar, Robert H. Mach, Robert B. Faryabi, Junwei Shi, Roger A. Greenberg

## Abstract

The response to Poly (ADP-ribose) polymerase inhibitors (PARPi) is dictated by homologous recombination (HR) DNA repair mechanisms and the abundance of lesions that trap PARP enzymes on chromatin. It remains unclear, however, if the established role of PARP in promoting chromatin accessibility impacts viability in these settings. Using a CRISPR based screen, we identify the PAR-binding Snf2-like ATPase, ALC1/CHD1L, as a key determinant of PARPi toxicity in HR-deficient cells. ALC1 loss reduced viability of BRCA mutant cells and enhanced their sensitivity to PARPi by up to 250-fold, while overcoming several known resistance mechanisms. ALC1 loss was not epistatic to other repair pathways that execute the PARPi response. Instead, ALC1 deficiency reduced chromatin accessibility concomitant with a decrease in the association of repair factors. This resulted in an accumulation of replication associated DNA damage and a reliance on HR. These findings establish PAR-dependent chromatin remodeling as a mechanistically distinct aspect of PARPi responses, implicating ALC1 inhibition as a new approach to overcome therapeutic resistance in HR-deficient cancers.

---

The complex chromatin environment of eukaryotic genomes necessitates rapid nucleosome remodeling events in response to specific cues. Such events initiate transcription at specific loci and are also required for the proper repair of DNA damage. Poly (ADP-ribose) polymerases, PARP1 and PARP2, are ideally suited to sense and transduce these events through their high affinity interactions with DNA lesions, which allosterically activate their enzymatic activity^1, 2^. PARP dependent histone PARylation promotes the rapid recruitment of PAR-binding effector proteins and mediates chromatin decompaction^3–6^. This common step in the recognition of a myriad of DNA lesions plays a key role in the early phases of several different repair pathways. These include single-strand break repair (SSBR), nucleotide excision repair (NER), and micro-homology mediated end joining (MMEJ), while more recently being implicated in classical non-homologous end joining (c-NHEJ) double-strand break repair and the protection of stalled replication forks^7–15^. Though PAR-binding by repair factors is a critical component of PARP responses to genotoxic stress, the extent to which PARylation directed chromatin remodeling impacts each of these repair pathways is not clearly understood.

Genetic inactivation of PARP1 becomes synthetically lethal in cells with loss of BRCA dependent homologous recombination (HR)^16, 17^. Given the multitude of PARP1 functions, the molecular basis of this genetic interaction remains unclear. However, the reliance of HR-deficient cells on PARP1 has been clinically exploited for the development of PARP inhibitors (PARPi)^18^. PARPi increase the number of genomic lesions that require BRCA dependent HR for repair in S-phase^19^. Additional PARPi cytotoxicity is dictated by its ability to trap PARP enzymes on chromatin^20^. While the high trapping ability of PARPi likely contributes to their clinical efficacy, this property is also proportional to the severity of side effects in HR proficient cells^21^. Another major caveat with the current PARPi regimen is the inevitable emergence of acquired resistance. Restoration of HR, reduction in PARP1 trapping, and drug efflux have been identified as mechanisms of resistance^22–24^. There are currently no approved means to effectively overcome PARPi resistance, making this an important limitation to their clinical efficacy.

Here we establish chromatin accessibility as a mechanistically distinct aspect of PARPi responses in HR-deficient cancers. Employing a CRISPR-Cas9 approach to target chromatin associated factors, we reveal the PAR-dependent nucleosome sliding enzyme, ALC1 (Amplified in Liver Cancer 1, also known as CHD1L), as a key determinant of PARPi toxicity in BRCA mutant cells. ALC1 deficiency caused synthetic sickness in combination with BRCA-mutation and conferred up to 250-fold increases in PARPi hypersensitivity in the context of HR-deficiency. Moreover, ALC1 loss was able to overcome several known PARPi resistance mechanisms as a consequence of this expanded therapeutic window. ALC1 function in the damage response was dependent on its ability to alter chromatin structure in a cooperative manner with PARP activity. These features establish PARP dependent chromatin accessibility as a new vulnerability in HR-deficient cancers.

## Results

### Loss of ALC1 confers PARPi hypersensitivity in BRCA-mutant cells

We performed a CRISPR-Cas9 genetic screen to identify loss-of-function mutations in chromatin regulators that generate hypersensitivity to the PARPi, olaparib, in BRCA mutated settings. The sgRNAs were designed to target the functional domains within proteins, an approach that has been shown to impart higher editing efficiency^25^. A CRISPR sgRNA library targeting 197 functional domains of 179 chromatin regulators was transduced into *Streptococcus pyogenes* SpCas9 expressing BRCA mutant cells. These included the ovarian and breast cancer cell lines UWB1.289 and SUM149PT, respectively, both of which contain mutations in exon 11 of *BRCA1* and express the hypomorphic BRCA1-11q isoform, and CAPAN-1, a pancreatic cancer line that expresses the non-functional 6174delT BRCA2 protein^26–28^. The screen was performed in the presence of 10 nM olaparib, which approximates the lethal dose 20 for these BRCA-mutant lines in a two-week clonogenic assay (Fig. 1a). Importantly, this low dose of olaparib allowed us to identify target genes that could enhance PARPi sensitivity across cell lines that harbor diverse mutations in BRCA1 and BRCA2 (Supplementary Table 1).

**Fig. 1.**
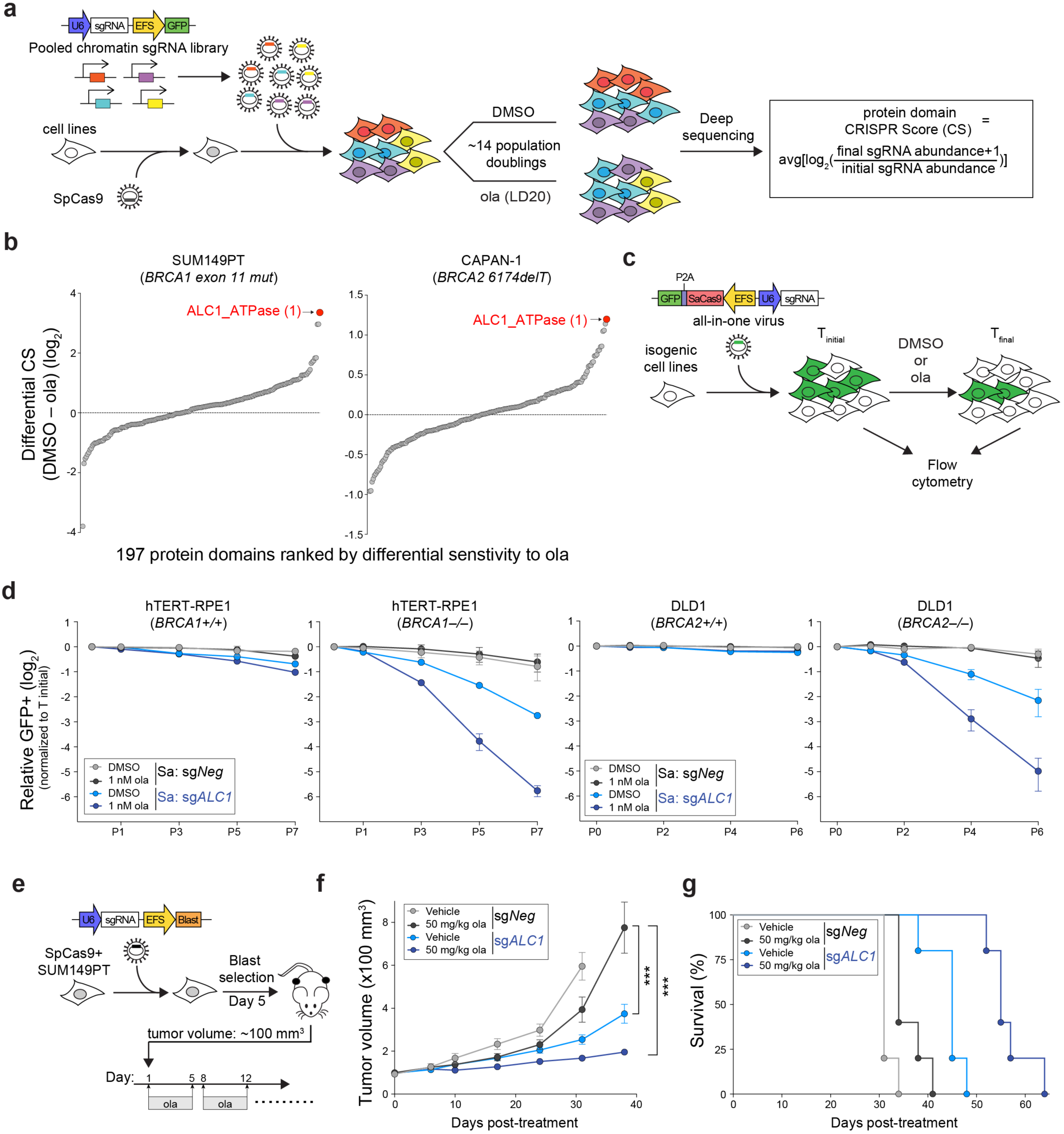
Loss of ALC1 reduces proliferation and confers olaparib hypersensitivity in BRCA-mutant cells. **a**, Schematic of the CRISPR screen to identify regulators of olaparib (ola) sensitivity. **b**, Protein domains ranked on the basis of CRISPR score (CS) for ola sensitivity in BRCA1 mutant SUM149PT cells (left) and BRCA2 mutant CAPAN-1 cells (right). **c**, Schematic of the GFP competition experiments. For a given cell line, T_initial_ indicates the day when maximum GFP expression is achieved for a sgRNA targeting an essential gene. T_final_ indicates the final day of the data collection. **d**, GFP competition assay in isogenic BRCA-mutant lines upon transduction of sg*Neg* or sg*ALC1* (n=3-6 independent transductions). Data are mean ± s.e.m., normalized to T_initial_. After every two population doublings, cells were passaged (P) and percent GFP was recorded. **e**, Schematic of the xenograft experiment. **f**, Measurement of tumor volume after ola treatment was initiated (n=10 tumors for sg*Neg* + ola, sg*ALC1* + vehicle, sg*ALC1* + ola and n=8 tumors for sg*Neg* + vehicle). Data are mean ± s.e.m. *P*-values are derived from a 2-way ANOVA. **g**, Kaplan-Meier survival analysis (n=5 per group). A significantly improved survival was observed in vehicle (light blue) and ola (dark blue) treated sg*ALC1* xenografts compared to the ola treated sg*Neg* counterpart (black), p=0.04 and p=0.0015 respectively using Log-rank (Mantel-Cox) test. \*\*\**P* ≤ 0.001. Cas9 from *S. aureus* (Sa) and *S. pyogenes* (Sp) was used as indicated.

ALC1 loss conferred olaparib hypersensitivity across all three cell lines and ranked as the top hit in both SUM149PT and CAPAN-1 cells (Fig. 1b, Extended Data Fig. 1a). By employing a growth-based competition assay that monitors changes in the percent of GFP, which is co-expressed from the sgRNA vector (Fig. 1c and Extended Data Fig. 1b), olaparib hypersensitivity was validated in all three BRCA-mutant lines using the seven different *ALC1* sgRNAs from the original CRISPR screen (Extended Data Fig. 1c). Loss of ALC1 also conferred a growth defect in all lines tested, albeit to a lesser degree in UWB1.289 cells, suggesting a synthetic sick relationship between ALC1 loss and BRCA-deficiency.

We next employed paired isogenic cell lines to examine if ALC1 loss selectively impacts the proliferation and PARPi sensitivity of *BRCA*-knockout (KO) cells in comparison to their *WT* counterparts. We utilized the *Staphylococcus aureus* SaCas9 system for ALC1 depletion to avoid competition with the SpCas9-*BRCA1* sgRNAs that were already present in one of the isogenic cell lines^29, 30^. Use of the dual SpCas9 and SaCas9 enzymes also allowed efficient simultaneous knockdown of two proteins in our subsequent experiments. Loss of ALC1 selectively impaired the proliferation of both *BRCA1* KO retinal pigmented epithelial cells (RPE1) and *BRCA2* KO colon cancer DLD1 cells, revealing a synthetic sick relationship with BRCA loss. Additionally, ALC1 depletion sensitized BRCA-mutant cells to low doses of olaparib (1 nM), while having minimal effects on their WT counterparts (Fig. 1d). This degree of sensitization upon ALC1 loss was striking given the *in vitro* olaparib half-maximal inhibitory concentration (IC_50_) of purified PARP1 enzyme is 10 nM^31^. This suggests that ALC1 loss hypersensitizes HR-deficient cells to PARPi concentrations that do not inhibit the majority of cellular PARP enzymatic activity. This selective hypersensitization of BRCA-deficient cells was corroborated in clonogenic survival and cellular ATP content-based viability assays that revealed up to ∼250-fold decreases in IC_50_ for olaparib upon ALC1 loss in several independent BRCA1- and BRCA2-mutant cell lines (Extended Data Fig. 2). The magnitude of this effect was larger in cancer cell lines compared to the primary hTERT-RPE1 cells, suggesting that ALC1 loss more strongly augments responses in cells with inherently higher levels of endogenous oxidative and replication stresses. ALC1 deficiency was also able to enhance sensitivity of BRCA2-mutant DLD1 cells to veliparib and talazoparib (Extended Data Fig. 3a). Together, these results demonstrate that ALC1 loss is detrimental to the proliferation of BRCA-mutant cells and decreases both the duration and dose of PARPi required for their selective killing.

**Fig. 2.**
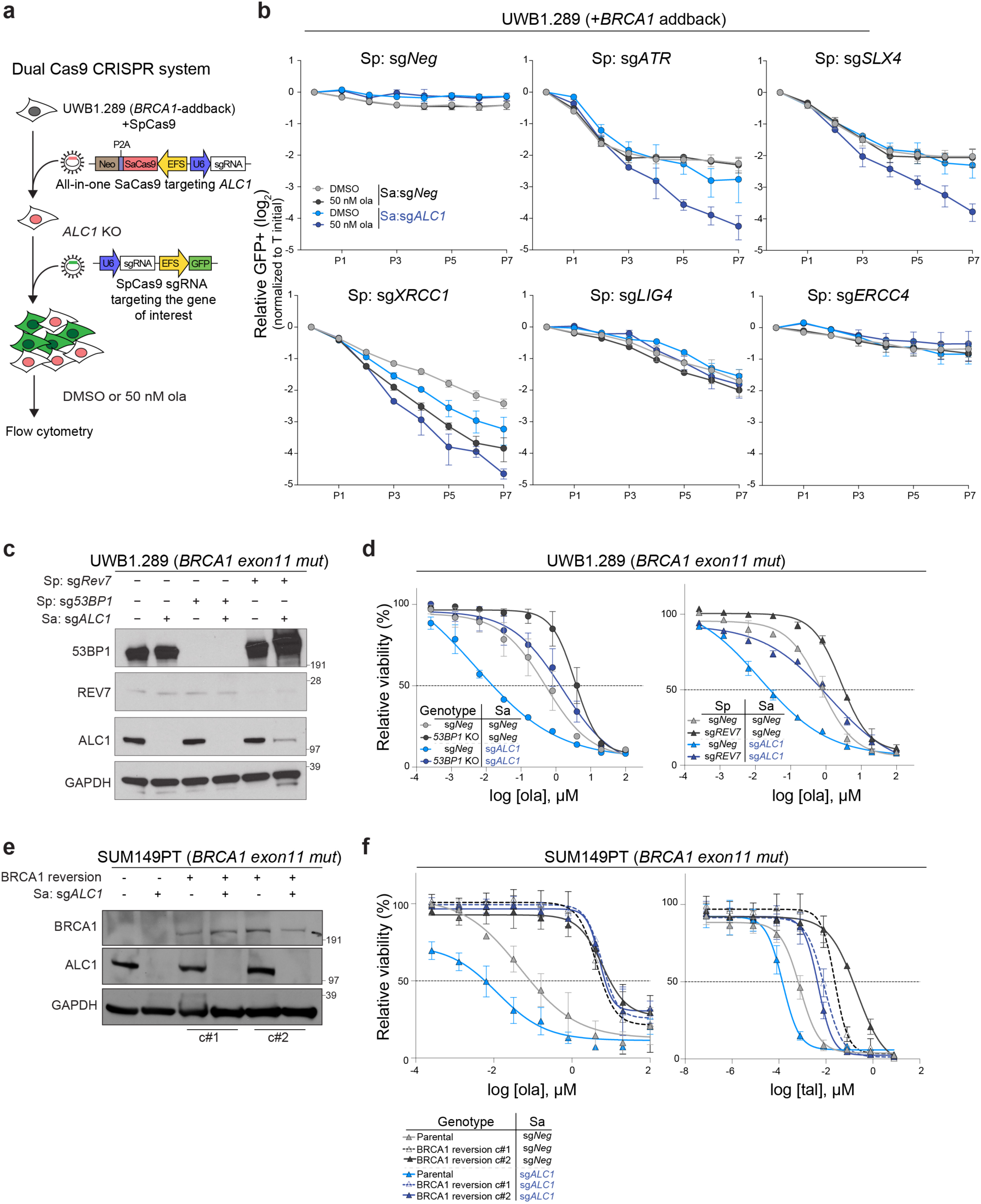
ALC1 loss causes PARPi hypersensitivity in HR-deficient cells. **a**, Schematic of the dual Cas9 CRISPR system for the GFP competition experiments. **b,** GFP competition experiment in the UWB1.289+*BRCA1* addback line to examine cell proliferation and ola sensitivity following the combined loss of ALC1 and the indicated DNA repair protein. Data are mean ± s.e.m., normalized to T_initial_. After every two population doublings, cells were passaged (P) and percent GFP was recorded (n=2-6 independent transductions). **c**, Immunoblot showing ALC1, 53BP1 and Rev7 depletion in UWB1.289 cells. **d**, Sensitivities of the indicated UWB1.289 cell lines to ola using the CellTiter-Glo assay; n=2 biologically independent experiments. **e**, Immunoblot showing BRCA1 and ALC1 levels in the indicated SUM149PT cells. **f,** Sensitivities of the indicated SUM149PT cell lines to ola (left) and talazoparib (tal) (right) using the CellTiter-Glo assay; n=2 biologically independent experiments. c#1 and c#2 indicates two different clones with restored BRCA1 reading frames. Data in **d** and **f** are mean ± range.

**Fig. 3.**
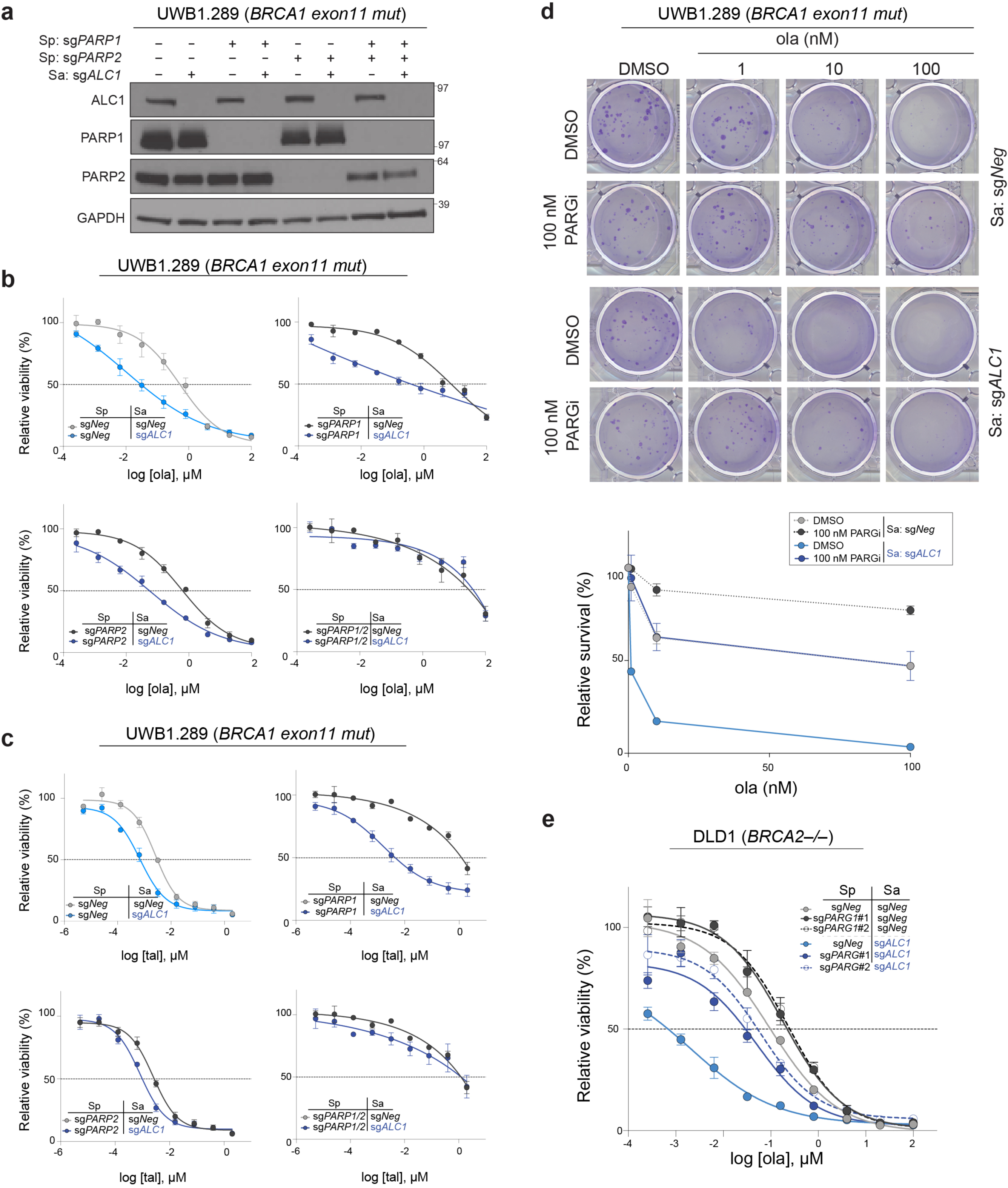
ALC1 loss mitigates PARPi resistance in BRCA-mutant cells that are deficient in PARP1 or PARG. **a**, Immunoblot showing ALC1, PARP1 and PARP2 depletion in UWB1.289 cells. **b-c**, Sensitivities of the indicated UWB1.289 cell lines to ola **(b)** and tal **(c)** using the CellTiter-Glo assay. Data are mean ± range from two biologically independent experiments. **d,** Representative images (top) and quantification (bottom) of the clonogenic survival assay of UWB1.289 cells treated with the indicated doses of ola and PARG inhibitor (PARGi). Colonies with more than 50 cells were included in the quantification. Data are mean ± range from two biologically independent experiments. **e,** Sensitivities of the indicated DLD1 *BRCA2*-/- cell lines to ola using the CellTiter-Glo assay. sg*PARG*#1 and sg*PARG*#2 indicate two sgRNAs targeting *PARG*. Data are mean ± s.e.m. from three biologically independent experiments.

We next examined if the proliferation defects and PARPi hypersensitivity observed upon ALC1 depletion in BRCA-mutant cells can be recapitulated in a tumor xenograft model. SUM149PT Cas9 cells were transduced with either sgNegative (sg*Neg*) or sg*ALC1* and the heterogeneous pool of engineered cells was subcutaneously implanted in NSG (NOD-*scid* gamma) mice on day 5 after transduction, when maximum sgRNA expression occurs. The time gap between sgRNA expression and complete protein depletion in the CRISPR editing system allowed the tumors to initially develop and reach ∼100 mm^3^, the stage when olaparib treatment was initiated (Fig. 1e). PARPi administration delayed growth in tumor volume compared to its vehicle treated counterpart. The rate of tumor growth in sg*ALC1* xenografts by itself was significantly slower compared to the olaparib treated sg*Neg* group, consistent with the synthetic sick interaction between ALC1 loss and BRCA mutation. Moreover, the volume of PARPi treated sg*Neg* tumors had increased by ∼7.8-fold compared to a ∼2-fold increase in sg*ALC1* counterparts by 38 days post administration of olaparib (Fig. 1f). Kaplan-Meier analyses indicated that the median overall survival was significantly longer for sg*ALC1* by itself compared to the olaparib administered sg*Neg* group and further extended in the PARPi-treated sg*ALC1* cohort (Fig. 1g).

Given that the xenografts were developed from a heterogeneous pool of sgRNA expressing cells, we examined if the eventual tumor growth in the sg*ALC1* group could be a consequence of the enrichment of cells that escaped genetic editing. Indeed, immuno-blot analysis of late stage tumors from sg*ALC1* group revealed the presence of residual ALC1 protein. In contrast, ALC1 protein was not detected at 15 days post transduction of sgALC1 in the cells that were used to inoculate the NSG mice (Extended Data Fig. 3b). These findings suggest that selective pressure exists to maintain ALC1 expression in BRCA mutant tumors that were able to proliferate in the presence of olaparib.

### PARPi sensitivity in ALC1 deficient cells is not epistatic with other DNA repair pathways

We systematically examined the genetic interaction between ALC1 and a selected group of repair pathways in relation to PARPi response. We designed an array of sgRNA libraries targeting DNA repair genes in the HR, NHEJ, SSBR, NER and MMEJ pathways. The proliferation of BRCA1-proficient UWB1.289 *ALC1* knockout (KO) cells was monitored following the loss of targeted genes in the absence or presence of 50 nM olaparib using the cell viability competition assay (Fig. 2a). Targeting ATR, BLM, SLX4, Mus81 and Fen1 sensitized *ALC1* KO cells to olaparib (50 nM). At this drug dose, no increase in olaparib sensitivity was observed in cells that express WT ALC1 (Fig. 2b, Extended Data Fig. 3c). ALC1 deficiency was also found to be synthetically sick with XRCC1 loss and further increased PARPi sensitivity, suggesting that ALC1 loss may generate lesions that increase reliance on canonical SSBR in addition to HR. Reciprocal experiments with ALC1 depletion in both sg*XRCC1* DLD1 cells and *XRCC1* KO hTERT-RPE1 cells also enhanced PARPi sensitivity. Notably, loss of ALC1 alone conferred a modest increase in talazoparib sensitivity to BRCA-proficient DLD1 cancer cells while having no effects in hTERT-RPE1 counterparts (Extended Data Fig. 4a-d). In contrast, ALC1 inactivation did not affect PARPi response following the loss of c-NHEJ or NER (Fig. 2b, Extended Data Fig. 3c). Collectively, these findings reveal that ALC1 loss is not epistatic to either of the primary determinants of PARPi response, HR or SSBR^32, 33^.

**Fig. 4.**
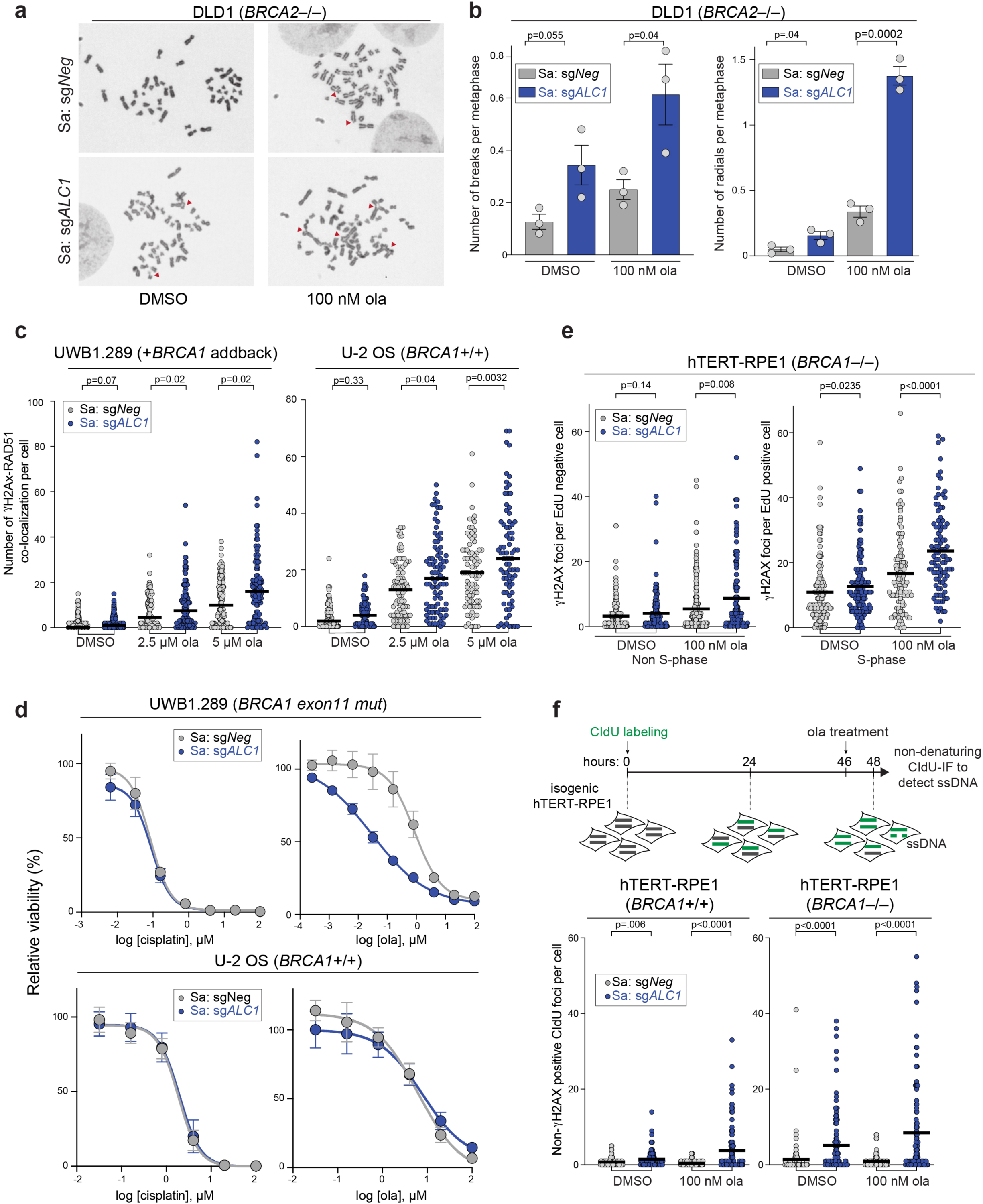
Loss of ALC1 increases genomic instability. **a-b**, Representative images **(a)** of the chromosomal aberrations (indicated by red arrow heads) and quantification **(b)** of breaks and radials per metaphase upon ALC1 depletion in DLD1 *BRCA2–/–* cells. Data are mean ± s.e.m. from three biologically independent experiments, *p*-value, unpaired *t*-test. For each experiment, at least 50 spreads were analyzed per sample. **c**, Quantification of γH2AX-RAD51 co-localization in indicated UWB1.289 + *BRCA1* add back and U-2 OS cells. Median is indicated. Data from two biologically independent experiments; *p* value, Mann-Whitney. For each experiment, at least 40 cells were quantified per sample. **d**, Sensitivities of the indicated cell lines to cisplatin and ola as determined by CellTiter-Glo assay. Data are mean ± s.e.m. from three biologically independent experiments. **e**, Quantification of γH2AX foci in non S-phase (left) and S-phase (right) cells. Median is indicated. Data from two biologically independent experiments; *p* value, Mann-Whitney. At least 50 cells were quantified per sample per experiment. **f**, Schematic and quantification of non-γH2AX positive CIdU foci in the indicated cell lines to specifically detect single strand (ss)-DNA that was not generated by end-resection at DSBs. Median is indicated. Data from two biologically independent experiments; *p* value, Mann-Whitney. For each experiment, at least 50 cells were quantified per sample. Cells were incubated with the indicated concentrations of ola for either 2 hrs (**f**) or 24 hrs (**b**, **c**, **e**).

### ALC1 deficiency restores PARPi sensitivity in cells with engineered resistance

The profound PARPi response in cells harboring combined ALC1 and BRCA deficiencies led us to examine if this relationship could be leveraged to overcome therapeutic resistance. Cell lines expressing hypomorphic BRCA1 become refractory to PARPi upon loss of 53BP1 or Rev7^34–37^. 53BP1 interacts with the Rev7-Shieldin complex to limit end resection and promote c-NHEJ. Depletion of either of these components permits partial restoration of HR in BRCA1-mutant cells coincident with both reduced radial chromosome formation and PARPi resistance^38–41^. Consistent with a partial effect on HR restoration, PARPi sensitivity and Rad51 foci formation ability of 53BP1 KO UWB1.289 cells was intermediate between the parental and BRCA1 addback counterparts (Extended Data Fig. 4e-f). ALC1 loss in either 53BP1 or Rev7 deficient BRCA1 mutant cells resulted in PARPi sensitivity (dark blue vs black), albeit at levels comparable to the parental control rather than the extreme sensitivity observed when ALC1 was targeted in 53BP1 or Rev7 WT UWB1.289 cells (light blue vs grey) (Fig. 2c, d).

This partial mitigation of PARPi hypersensitivity in the absence of 53BP1 or REV7 suggests that ALC1 deficiency in BRCA1 mutant cells generates lesions that are subjected to toxic c-NHEJ. In accordance, ALC1 deficient UWB1.289 cells formed increased radial chromosomes upon PARPi treatment that were eliminated by concomitant 53BP1 depletion (Extended Data Fig. 4g-h). Conversely, increased chromosome breaks and Rad51 foci were observed in PARPi treated ALC1 deficient *53BP1* KO BRCA1 mutant cells. This may account for the continued sensitivity of ALC1 deficient *53BP1* KO UWB1.289 cells, consistent with reports that accumulation of genomic breaks is an underlying cause of PARPi cytotoxicity in HR-proficient cells^29, 33^.

To examine if complete HR restoration would negate the impact of ALC1 on PARPi response, we extended this analysis to reversion mutations that produce full length BRCA1 protein. ALC1 loss failed to cause olaparib sensitivity in SUM149PT cells that have become resistant owing to the restoration of the BRCA1 reading frame^42^. In contrast, some degree of PARPi sensitivity was observed in ALC1 deficient, *BRCA1* revertant cells treated with talazoparib (Fig. 2e, f). We posit that despite restoration of HR, the more potent PARP trapper, talazoparib, may produce some degree of toxicity due to the nature of genomic lesions that accumulate in ALC1 deficient cells (Extended Data Fig. 4a-d).

Trapping of PARP1 on chromatin by PARPi is thought to contribute to PARPi cytotoxicity in BRCA-mutant cells^20^. Reduced expression of PARP1 is therefore associated with PARPi resistance^43, 44^. Remarkably, ALC1 loss was able to enhance PARPi sensitivity of PARP1 depleted BRCA1-mutant cells. At lower PARPi concentrations, cells depleted of both PARP1 and ALC1 were more sensitive to either olaparib or talazoparib compared to parental controls (Fig. 3a-c). Given that PARPi also traps PARP2, this could account for the continued PARPi sensitivities in ALC1 and PARP1 depleted UWB1.289 cells^20^. Using a quantitative immunofluorescence assay, we observed increased PARP2 trapping in ALC1 deficient cells compared to the parental control (Extended Data Fig. 5a)^45^. Strikingly, the combined loss of PARP1 and 2 rendered ALC1 deficient BRCA1 mutant cells fully resistant to PARPi (Fig. 3a-c). We propose that the abundance of genomic lesions in ALC1 deficient cells permits increased trapping of both PARP1 and PARP2. This accentuated PARP2 trapping in ALC1 null cells accounts for the continued efficacy of PARPi in the absence of PARP1.

**Fig. 5.**
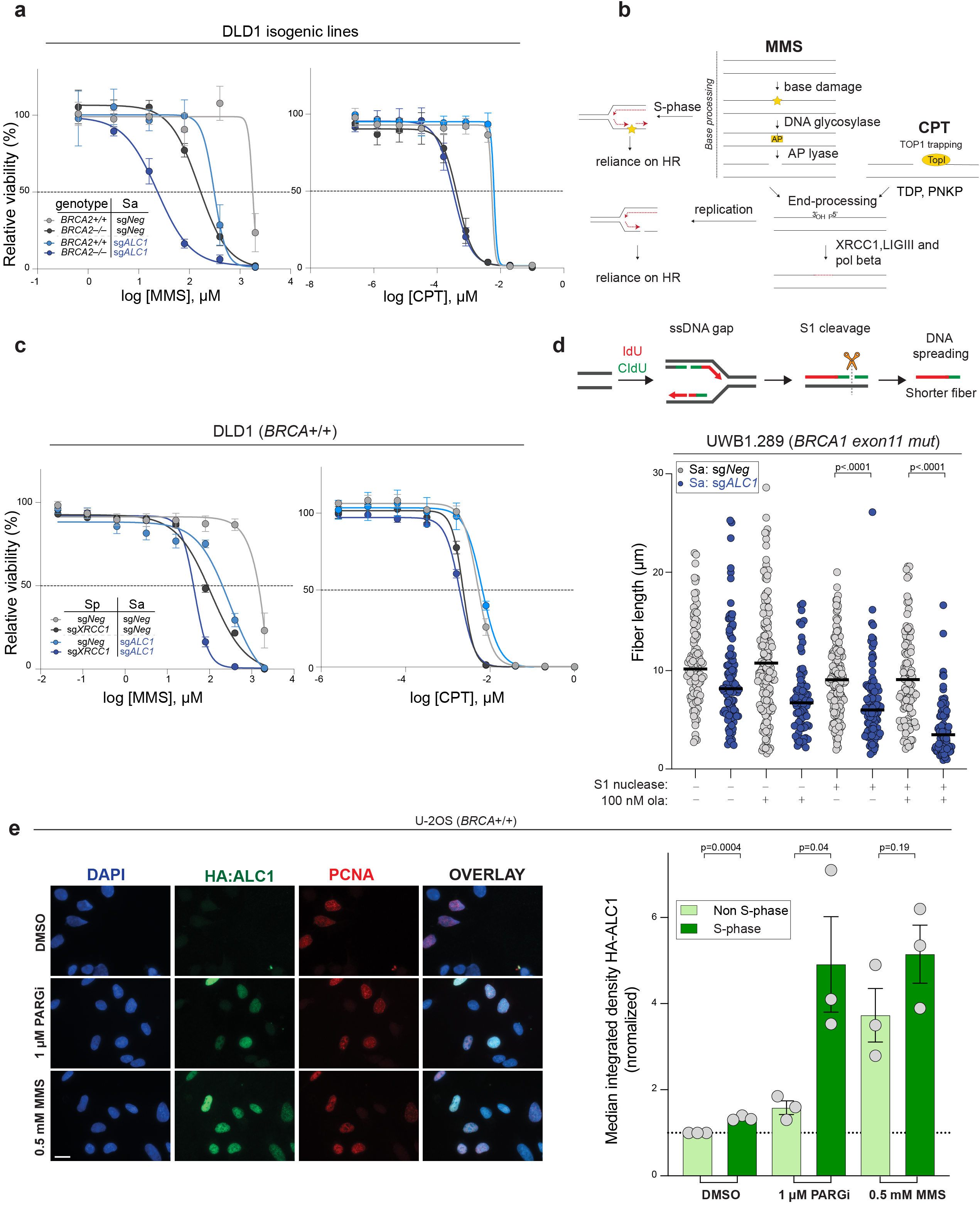
ALC1 function in base damage repair is not epistatic with HR and single-strand break repair (SSBR). **a**, Sensitivities of the indicated DLD1 isogenic lines to methyl methanesulfonate (MMS) and camptothecin (CPT) in the CellTiter-Glo assay. Data are mean ± s.e.m. from three biologically independent experiments. **b,** Schematic of SSBR pathway. While base damage requires additional processing events at the chromatin, CPT traps TOP1 on already nicked substrates. Replication of damaged bases results in gaps and SSBs, that get converted to DSBs and increase reliance on HR for repair. **c,** Sensitivities of the indicated DLD1 cell lines to MMS and CPT in the CellTiter-Glo assay. Data are mean ± s.e.m. from three biologically independent experiments. **d,** Schematic and quantification of fiber length in the absence and presence of S1 nuclease in the indicated cell lines. Median is indicated. Data from two biologically independent experiments. For each experiment, at least 45 fibers were quantified per sample. Cells were incubated with 100 nM ola for 24 hrs. *P* value, Mann-Whitney. **e,** Representative images (left) and quantification (right) of HA-ALC1 localization to chromatin upon the indicated treatments. Scale bar, 50 microns. The median value was normalized to non S-phase untreated control. Data are mean ± s.e.m. from three biologically independent experiments, *p* value, unpaired Student’s *t*-test. For each experiment, at least 50 cells were analyzed per sample.

Loss of poly(ADP-ribose) glycohydrolase (PARG) activity represents another PARPi resistance mechanism in BRCA2-deleted murine tumor-derived cells^46^. Accumulation of PAR in the absence of PARG activity correlates with less PARP trapping and resistance. Loss of ALC1 was able to enhance PARPi sensitivity in cells that became resistant either owing to chemical inhibition of PARG or its genetic depletion, albeit some degree of resistance was still observed compared to ALC1 loss alone (Fig. 3d-e). Collectively, these observations (Figs. 2 and 3) demonstrate that ALC1 loss restores PARPi efficacy across a broad range of established resistance mechanisms.

### ALC1 deficiency increases genomic instability and reliance on BRCA-dependent HR

We next characterized the nature of the genomic lesions that arose from ALC1 loss in PARPi treated BRCA-mutant. Most experiments were performed at 100 nM olaparib, a concentration that did not affect the viability of BRCA-WT cells that lack ALC1 in a two-week viability assay (Extended Data Fig. 2b). Depletion of ALC1 in DLD1 BRCA2-deficient cells increased breaks and radial chromosomes, which were exacerbated by olaparib treatment (Fig. 4a, b). Loss of ALC1 increased Rad51 foci formation upon olaparib treatment in BRCA-proficient U-2 OS and UWB1.289 cells, suggesting the genesis of genomic lesions that require BRCA-dependent HR (Fig. 4c and Extended Data Fig. 5b). ALC1 loss did not enhance sensitivity to cisplatin in BRCA-proficient or deficient settings, emphasizing that it does not generally contribute to homology-directed repair as is commonly observed for other factors that affect PARPi sensitivity through HR or MMEJ (Fig. 4d) ^32, 47^.

ALC1 depletion in BRCA-mutant cells increased γH2AX foci specifically in S-phase, which were further enhanced upon PARPi treatment, consistent with the generation of replication-coupled DSBs (Fig. 4e and Extended Data Fig. 5c). We next quantified single strand-DNA (ss-DNA) in cells using non-denaturing CIdU immunofluorescence. ALC1 loss alone resulted in a significant increase in the number of non-γH2AX positive CIdU foci, independent of BRCA status, albeit the magnitude was higher in BRCA1-mutant cells. Non-γH2AX positive CIdU foci in ALC1 deficient cells were further elevated upon olaparib treatment in both cases (Fig. 4f).

ALC1 deficiency has been reported to confer sensitivity to agents that induce single-strand breaks (SSBs), albeit some discrepancies exist between studies regarding the response to different agents^48–50^. We were able to recapitulate the reported MMS hypersensitivity across different ALC1 depleted cell lines, with a notable exception in hTERT-RPE1 cells. BRCA loss exacerbated MMS sensitivity in ALC1 depleted cells consistent with channeling of unrepaired SSBs into HR during DNA replication^11, 51^. In contrast, ALC1 depletion did not confer sensitivity to camptothecin (CPT) (Fig. 5a, Extended Data Fig. 6a-b), revealing that ALC1 is not generally required for SSBR. While both MMS and CPT increased SSBs and induced PAR synthesis, the extent of chromatin remodeling required to repair genomic lesions arising from these agents may differ. MMS introduces base methylation and requires complex lesion processing events that utilize BER and HR. On the other hand, CPT traps the TOP1 enzyme on already nicked DNA^52^. Thus, while repair in response to MMS would require nucleosome sliding to provide accessibility to the damaged base, the same extent of remodeling may not be required for repair of CPT induced lesions (Fig. 5b). In support of this possibility, ALC1 has been shown to function in the pre-incision step of NER by recognizing base damage in a PARP1 dependent manner^14^.

**Fig. 6.**
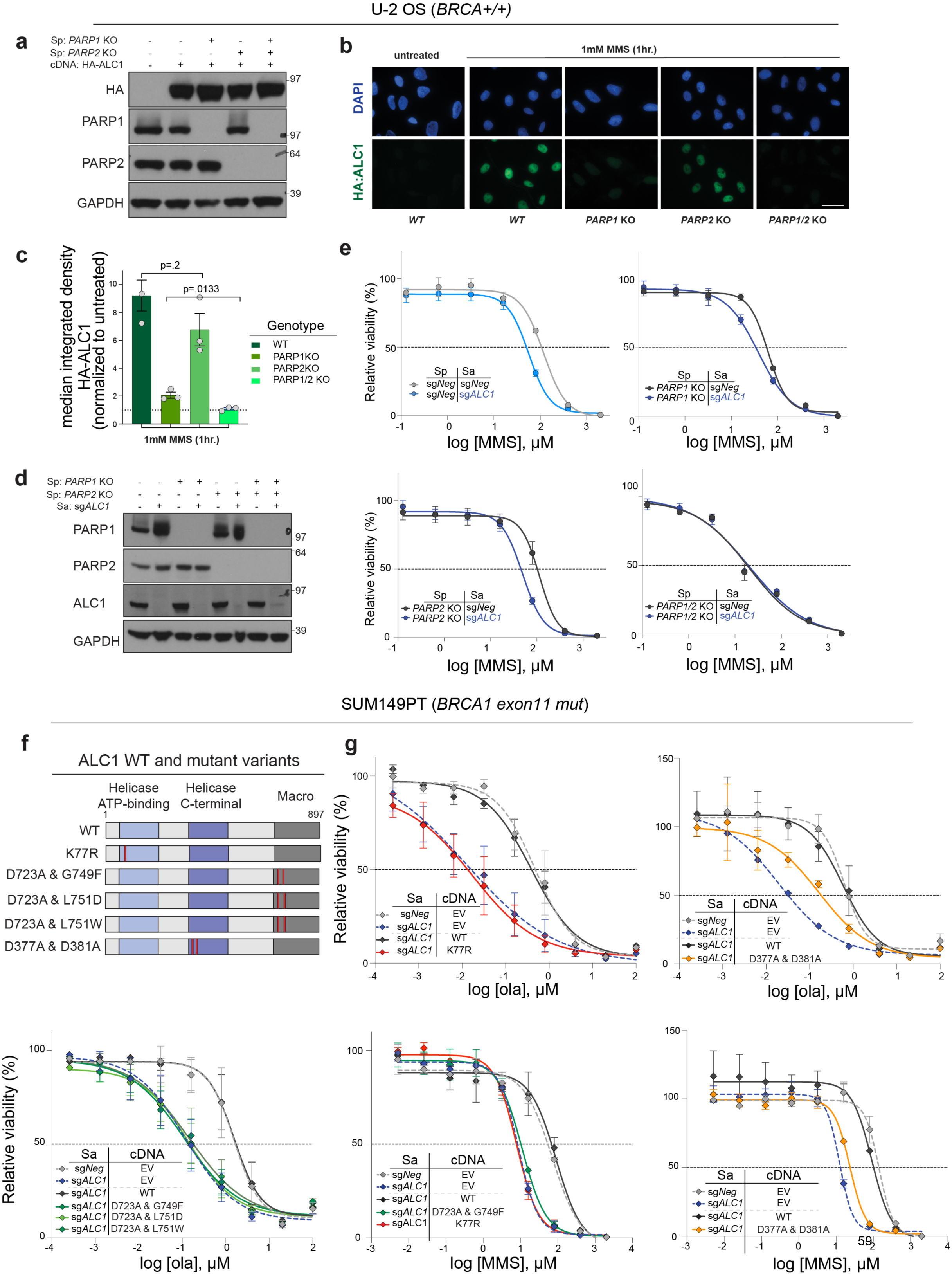
The ATPase and macro domains of ALC1 are essential to prevent replication-associated damage in response to PARPi and MMS. **a**, Immunoblot showing expression of HA-ALC1 in indicated cell lines. **b-c**, Representative images (**b**) and quantification (**c**) of HA-ALC1 localization to chromatin, scale bar 50 microns. For each cell line, the median value upon MMS treatment was normalized to its respective untreated control. Data are mean ± s.e.m. from three biologically independent experiments, *p* value, unpaired Student’s *t*-test. For each experiment, at least 50 cells were analyzed per sample. **d**, Immunoblot showing depletion of ALC1 in the indicated cell lines. **e**, Sensitivities of the indicated U-2 OS lines to MMS in the CellTiter-Glo assay. Data are mean ± s.e.m. from three biologically independent experiments. **f**, Domain organization and mutants of ALC1 used in this study. The helicase ATP-binding (light blue), helicase C-terminal (dark blue), and macro domains (dark grey) are indicated. Red bars show the position of the residues that were mutated. **g**, Sensitivities of SUM149PT cells expressing various ALC1 mutants to ola and MMS using the CellTiter-Glo assay (EV: empty vector and WT: wild type). Data are mean ± range from two biologically independent experiments.

ALC1 loss specifically enhanced MMS and not CPT sensitivity in XRCC1 depleted cells (Fig. 5c and Extended Data Fig. 6b). Based on these findings we propose that ALC1 acts upstream of XRCC1 and HR in repairing alkylation damage to promote the repair of base damage. The presence of persistent chromatin lesions may lead to the generation of replication-coupled ss-gaps^53^. Indeed, loss of ALC1 increased the S1 nuclease sensitivity of nascent replication tracts and this phenotype was exacerbated upon PARPi treatment (Fig. 5d and Extended Data Fig. 6c). Together, these results demonstrate that ALC1 prevents the accumulation of toxic genomic lesions that feed into SSB and HR repair mechanisms.

### ALC1 localizes to chromatin in a PARP1 and PARP2 dependent manner

ALC1 recruitment to damaged chromatin is reliant on PARylation ^48, 54–56^. It is therefore intriguing that loss of this PAR-binding protein increases PARPi sensitivity. We reasoned that owing to its nanomolar binding affinity for PAR chains, ALC1 localization to damaged chromatin would not be diminished by low dose or short duration PARPi treatments that do not recapitulate combined loss of PARP1 and PARP2^57^.

ALC1 association with chromatin was quantified using immunofluorescence on detergent pre-extracted cells. Treatment of cells with PARGi revealed accumulation of ALC1 specifically on S-phase chromatin. This suggests that in the absence of damaging agents, ALC1 primarily associates with replicating DNA, consistent with reports that PAR levels are highest in S-phase^58^. ALC1 could be detected in both S- and non-S-phase cells following MMS treatment and did not require PARGi to elicit detectable association (Fig. 5e). MMS-induced ALC1 chromatin localization was retained in cells pre-incubated with 100 nM olaparib for 24 hrs or after 4 hours treatment with 5 µM olaparib (Extended Data Fig. 7a-c). Notably, a requirement of high PARPi concentration to abolish the recruitment of PAR binding proteins has been reported. For example, a concentration of 10 µM olaparib was required to completely nullify recruitment of the PAR-binding protein XRCC1 to damaged chromatin^57^.

**Fig. 7.**
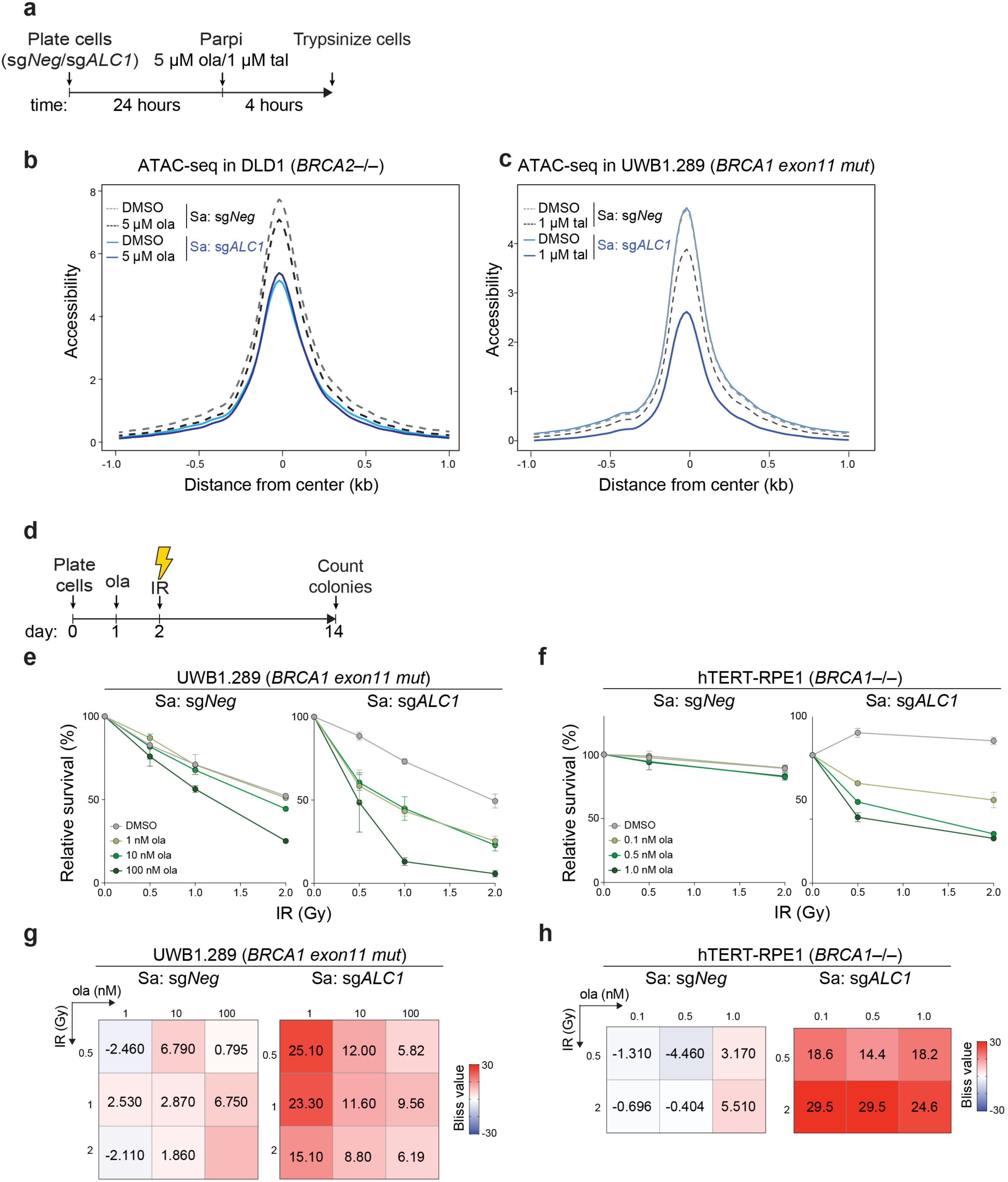
ALC1 and PARP activities cooperate to promote chromatin accessibility during the DNA damage response. **a**, Schematic of the experiment used to examine the effects of ALC1 loss and PARPi treatment on chromatin accessibility **b,** ATAC-seq analysis in DLD1 *BRCA2-/-* cells to assess global accessibility of chromatin. Cells were treated with ola for 4 hours. Data is from three replicates for each condition. **c,** ATAC-seq analysis in UWB1.289 cells to assess global accessibility of chromatin. Cells were treated with tal for 4 hrs. Data is from three replicates of untreated sg*Neg* and sg*ALC1*+tal and two replicates of sg*Neg*+tal and untreated sg*ALC1* cells. For plotting of both the ATAC-seq graphs **(b,c)** the 2000 base-pairs flanking the center of the accessible sites (i.e. from −1 to 1 on x-axis) were divided into equally sized 20 bp regions. The differential accessibility was calculated at each of these 20 bp regions. **d,** Schematic of the experiment to examine the effects of combining low doses of PARPi with ionizing radiation (IR) in ALC1 depleted cells. **e-h,** Quantification of clonogenic survival **(e,f)** and heat map of bliss scores **(g,h)** obtained from UWB1.289 and *BRCA1–/–* hTERT-RPE1 cells treated with the indicated doses of ola and IR. Data are mean ± range from two biologically independent experiments. Bliss score >0, synergistic; Bliss score <0, antagonistic; Bliss score = 0, additive. Number of colonies in IR-treated conditions were normalized to their respective un-irradiated counterparts. Colonies with more than 50 cells were included in the analysis.

To determine if ALC1 chromatin association is completely PAR dependent, we examined the effects of genetically depleting both PARP1 and PARP2 enzymes. PARP1 loss resulted in approximately 80% reduction in ALC1 localization to damaged chromatin as assessed by quantitative immunofluorescence. PARP2 partially compensated for the absence of PARP1. ALC1 recruitment to chromatin was absent in cells with the combined loss of both PARP1 and PARP2 (Fig. 6a-c). In accordance, ALC1 loss did not further increase the MMS sensitivity of *PARP1* and *PARP2* double KO cells (Fig. 6d-e).

### ALC1 responses to PARPi and MMS require its chromatin remodeling and PAR binding activities

We investigated the relevant biochemical properties underlying ALC1 function to PARPi responses in HR deficient cells. ALC1 is a bona fide member of SNF2 superfamily, whose members are characterized by seven signature motifs ^59^. The ALC1 N-terminus ATPase domain has 64% similarity to the SNF2 like chromatin remodeling enzyme, SMARCA1. Both ALC1 and SMARCA1 share similar nucleosome re-positioning properties *in vitro* that depend on their respective ATPase activities^48, 54, 60^. The K77R mutation in the Walker A motif of the ALC1 ATPase domain abrogates nucleosome sliding activity. Notably, overexpression of the K77R ALC1 mutant led to a pronounced loss of viability of BRCA-mutant RPE1 and SUM149PT cells, suggesting a dominant negative phenotype. Depletion of the endogenous protein in the surviving population of K77R expressing cells resulted in PARPi hypersensitivity comparable to ALC1 deficient cells (Fig. 6f, g and Extended Data Fig. 8a,b).

**Fig. 8.**
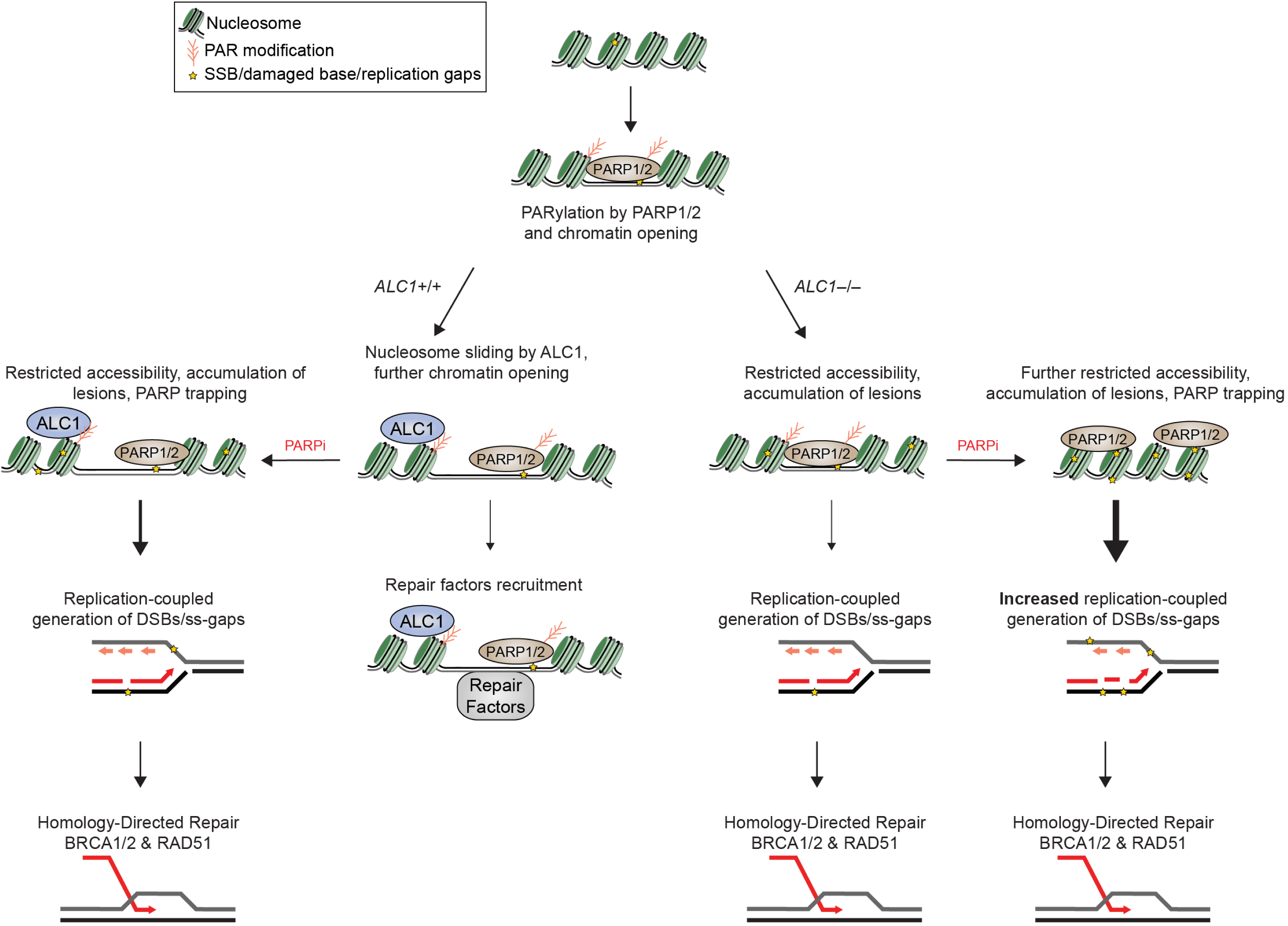
Model depicting cooperation between ALC1 and PARylation in response to DNA damage. PARylation results in chromatin decondensation and ALC1 recruitment. Nucleosome sliding by ALC1 further enhances chromatin accessibility. Combined loss of ALC1 and PARP activities reduces the access of base damage repair factors to the chromatin. Unrepaired base and single strand breaks (SSBs) result in the generation of replication coupled ss-gaps and DSBs during S-phase that necessitate repair by BRCA-dependent HR.

Like other chromatin remodelers, the basic histone H4 tail is essential for ALC1 nucleosome sliding activity ^48^. We mapped the acidic residues in the ALC1 ATPase domain that would interact with the basic histone H4 tail based on homology to a previously reported cryo-EM structure of the ATPase domain of an ISWI remodeler (PDB:6PWF)^61^. These H4 interaction residues (D377, D381) are conserved across multiple families of chromatin remodelers (Extended Data Fig. 8c). Mutation of these acidic residues to alanine reduced the association of ALC1 with histone H4 (Extended Data Fig. 8d). D377A+ D381A mutations in ALC1 failed to rescue PARPi responses in BRCA1 mutant SUM149PT cells (Fig. 6 f,g).

To examine the contribution of the macrodomain, we tested a previously reported D723A substitution that reduces binding affinity for PAR^48^. Complementation with the D723A mutant still rescued PARPi sensitivity in ALC1-deficient BRCA1 mutant cells (Extended Data Fig. 8a). This may reflect the ability of ALC1 to localize to the damage sites with reduced PAR levels^55^. To abrogate the residual PAR-binding and damage localization activity of the D723A mutant, other points mutations were rationally designed based on the available crystal structure of the PAR-binding module of AF1521 (PDB:2BFQ)^62^. Amino acids G749 and L751 in ALC1 structurally correspond to AF1521 G41 and V43 residues, which form hydrogen bonding interactions with the ADP-ribose in the macrodomain. While cellular complementation with the single mutants D723A, G749F, L751D and L751W showed complete or partial rescue of PARPi sensitivity, no protection was observed with any of the double mutants (D723A+G749F, D723A+L751D, D723A+L751W) (Fig. 6f, g and Extended Fig. 8e). The ATPase activity, H4 interaction and macro domain were also essential for the rescue of MMS sensitivity in ALC1 depleted cells (Fig. 6f, g). Together these results demonstrate that both PAR recognition and nucleosome sliding by ALC1 is essential for its ability to protect BRCA-mutant cells from PARPi and alkylation damage.

### ALC1 and PARP activity cooperate to promote chromatin accessibility

Reliance on PAR binding and nucleosome sliding activities for response to PARPi and MMS suggests ALC1 involvement in PARP dependent chromatin remodeling. ALC1 has been reported to affect chromatin decondensation at laser-induced damage sites ^63^. To examine if ALC1 loss alone or in combination with PARPi altered genome-wide chromatin accessibility we performed ATAC-seq (Assay for Transpose-Accessible Chromatin) analysis in BRCA2-mutant DLD1 cells. The interaction between ALC1 and newly synthetized PAR for altering chromatin accessibility was assessed in cells treated with 5 µM olaparib for four hours (Fig. 7a). A longer duration of treatment was avoided to negate chromatin changes owing to differences in cell cycle. Pairwise comparisons of accessible sites using DESeq2 analysis did not identify large locus-specific changes upon PARPi treatment, ALC1 depletion alone or in combination with PARPi (log2 fold change ≥0.5 or ≤-0.5, FDR <0.05) (Supplementary Table 2). The lack of locus specific changes suggested that ALC1 does not play a major role in regulating gene expression. We corroborated this conclusion by performing RNA-seq analysis in ALC1 depleted *delp53 BRCA1-/-* hTERT-RPE1 and *BRCA2-/-* DLD1 cells (Supplementary Table 3). No consistent gene expression changes were observed across the two cell lines.

We next examined if ALC1 loss had an effect on overall chromatin accessibility. Metagene analysis revealed a significant 1.6 fold overall reduction in chromatin accessibility upon ALC1 depletion alone. A four-hour olaparib treatment resulted in 1.3 and 1.84 fold changes in overall accessibility in sg*Neg* and sg*ALC1* cell respectively compared to DMSO treated control (paired t-test p-value <1E-15) (Fig. 7b). We reasoned that for a short duration treatment, a more potent PARPi may manifest in larger accessibility changes. To examine this possibility, we performed ATAC-seq analysis in UWB1.289 cells, which are less sensitive to ALC1 loss in the absence of drug treatment and thus allowed us to examine PARPi responses without the confounding issues of toxicity from purely genetic interactions. ALC1 loss alone revealed no significant reduction in overall chromatin accessibility in UWB1.289 cells. In contrast, ALC1 depletion had a profound impact on chromatin accessibility when combined with the potent PARPi, talazoparib. Compared to the untreated control, a four hour administration of talazoparib (1 µM) alone resulted in a 1.3 fold overall reduction in chromatin accessibility (paired t-test p-value < 1E-15) while combining talazoparib with ALC1 loss yielded a 3.5 fold reduction (paired t-test p-value < 1E-15) (Fig. 7c). ALC1 remained bound to chromatin during these PARPi treatments (Extended Data Fig. 7, 9a-b). These data reveal that the concerted activities of ALC1 and PARP regulate chromatin accessibility.

Chromatin decondensation facilitates accessibility of DNA repair proteins to genomic lesions^64^. We therefore performed chromatin fractionation studies to probe for the association of various DNA repair factors. Strikingly, loss of ALC1 alone in DLD1 BRCA2-mutant cells reduced the association of several proteins dedicated to processing base damage (Extended Data Fig. 9a,c). This included the DNA glycosylase NTHL1, the apurinic/apyrimidinic endonuclease APE1, and XRCC1. These reductions were more pronounced upon combining ALC1 loss with olaparib, suggesting that ALC1 and PARP activities act together to ensure chromatin accessibility to certain repair factors. In contrast, no effects were observed on the association of other repair or PAR-binding proteins, such as 53BP1, CHD4 or PARG. Similar changes in chromatin association of repair factors after PARPi treatment were also observed in UWB1.289 cells (Extended Data Fig. 9b). XRCC1 levels on chromatin were significantly lower in PARPi treated ALC1 deficient UWB1.289 cells compared to its WT counterpart, even in the presence of damaging agents that induce PARylation (Extended Data Fig. 9d-g).

Given the involvement of PARP-dependent responses to a host of DNA lesions, we posited that ALC1 functions cooperatively with PARylation to mediate a broader range of DNA damage responses. To test this hypothesis, we examined if combining PARPi with ALC1 depletion affects responses to other genotoxic insults. Clonogenic survival of BRCA1-mutant hTERT-RPE1 or UWB1.289 cells were assessed following ionizing radiation (IR) at 24 hrs after PARPi treatment. ALC1 loss by itself did not increase sensitivity to IR in either cell type. In contrast, treatment with nanomolar concentrations of PARPi showed synergy with ALC1 loss to achieve cell killing at low doses of IR in both BRCA1-mutant cell lines. Only modest or no synergy with IR was observed at these PARPi concentrations in ALC1 proficient cells (Fig. 7d-h, Extended Data Fig. 10). These results lend support to a role for ALC1 in promoting diverse PARP dependent damage responses.

## Discussion

This study identifies ALC1 as a new vulnerability in HR deficient cancers. ALC1 deficiency reduced the proliferation of BRCA-mutant cells and enhanced the efficacy of PARPi in a mechanistically distinct manner from perturbations in canonical DNA repair pathways. ALC1 was not required for SSBR or HR, the known determinants of PARPi sensitivity^29, 32, 33, 65–67^. Instead, ALC1 modulates chromatin to enable processing of complex lesions such as those arising from base alkylation damage. Combined loss of ALC1 and PARP activities reduced access to repair factors that process base damage. We propose that PARPi treatment in ALC1 deficient cells results in an accumulation of persistent lesions that necessitate a transition to BRCA-dependent HR repair during S-phase (Fig. 8). The non-epistasic interaction between HR and ALC1 for PARPi response and the increased accumulation of replication-coupled DSBs and ss-DNA gaps in ALC1 deficient BRCA-mutant cells lend support to this model.

PAR-dependent changes in chromatin accessibility were reported nearly four decades ago, yet its relationship to the selective toxicity of PARPi in HR deficient cells has remained largely unexplored^4^. The reductions in genome-wide chromatin accessibility upon ALC1 loss in BRCA-mutant cells did not associate with any specific locus or transcriptional processes. This is suggestive that levels of endogenous damage may dictate reliance on this remodeler. Synthetic sickness with ALC1 loss in untreated BRCA-mutant cancer cells and tumor xenografts is consistent with this hypothesis. Moreover, requirements for PAR recognition, histone H4 interaction, and ATPase activity directly connect ALC1 driven nucleosome sliding to PARPi responses in BRCA mutant cells.

ALC1 chromatin localization and function in the damage response was completely dependent on the combined activities of PARP1 and PARP2. The nanomolar binding affinity of the ALC1 macrodomain to PAR chains may allow ALC1 to access reduced levels of chromatin PARylation in PARP1 deficient settings and upon low doses or short durations of high dose PARPi treatment performed in our studies^55^. Interestingly, ALC1 has also been implicated in protecting PARylated proteins from PARG hydrolysis, which may further permit it to modulate PAR-dependent chromatin accessibility^68^. These findings address the paradox of how loss of a PAR binding protein can enhance PARPi sensitivity.

We hypothesize that genomic aberrations that arise in the absence of ALC1 trap a subset of inhibited PARP1 and PARP2 enzymes. Collectively, these events make an important contribution to the extreme PARPi sensitivity observed in BRCA mutant cells that harbor ALC1 deficiency. The requirements for both PARP1 and PARP2 loss for resistance to PARPi or MMS provides genetic proof that PARP2 also becomes important in the setting of ALC1 loss. This differentiates PARPi response determinants from BRCA deficiency alone, where inhibition and trapping of PARP1 is thought to be the major PARP enzyme responsible for therapeutic window^20, 44^.

The selective sensitivity of ALC1 deficient cells to alkylating damage indicates that ALC1 directed nucleosome sliding plays a vital role specifically in DNA repair processes that require extensive chromatin remodeling, rather than to all PARP dependent events. Moreover, reliance of ALC1 null cells on HR and SSB repair points to a role in preventing the generation of DNA lesions that channel into these pathways. Persistent base lesions in ALC1 deficient cells may impede replication fork progression and increase the frequency of replication-coupled gaps that trap PARPi. Interestingly, several recent studies reported the accumulation of ss-DNA enriched post-replicative territories upon treatments with alkylating agents or PARPi ^69, 70^. Replication restart by re-priming events downstream of DNA lesion can leave ss-DNA gaps to be filled post replication^71, 72^. The gaps could be filled in by translesion polymerases or repaired by replication restart or template switching with the sister chromatid ^73^. The role of HR proteins in replication restart and promoting template switch may account for the lower frequency of gaps in BRCA-proficient settings upon ALC1 loss (Fig.4f)^74, 75^. Though ALC1 responds to PAR-inducing MMS damage irrespective of cell cycle, it appears to predominantly act during S-phase in unperturbed cells when PARylation is highest (Fig. 5e). In agreement, ALC1 has been reported as a component of active replisomes, raising possibilities for a role in replication associated repair^76^. The presence of a PAR dependent nucleosome slider at the replisome may ensure efficient processing of chromatin lesions and progression of replication forks without the formation of ss-DNA gaps.

Based on the ability of ALC1 to promote the recruitment of multiple BER factors to chromatin and its non-epistasis with XRCC1 for both PARPi and MMS responses, we propose that accumulation of damaged bases upon ALC1 loss is responsible for the generation of replication gaps during S-phase. Interestingly, data from a recent genome-wide CRISPR screen demonstrated that loss of enzymes involved in processing base damage confers PARPi sensitivity^77^. In combination with our findings, these observations suggest that increased accumulation of base damage and the resultant genesis of replication-coupled gaps may provide a new avenue for enhancing PARPi cytotoxicity. ALC1 loss has also been reported to produce PARPi sensitivity in genome wide screens in BRCA-proficient cell lines passaged for over several weeks at olaparib concentrations of 0.5-2 μM (50-200 fold higher doses than used in our studies) or 5 µM veliparib, albeit investigation of underlying mechanisms was not described in these studies ^29, 77, 78^. These findings, together with our data from isogenic cell lines (Extended Data Fig. 2) indicate that a large therapeutic window can be obtained by targeting ALC1 in tumors with HR deficiency in comparison to HR proficient normal tissues.

Development of ALC1 inhibitors could represent a new approach to address the clinical problem of resistance to PARPi. Other determinants of PARP inhibitor sensitivity typically affect responses to genotoxic agents that necessitate HR repair. For example, sensitivity to platinum chemotherapy is a positive predictor of PARP inhibitor clinical efficacy and an FDA mandated indication for the use of PARP inhibitors in ovarian cancer. Nonetheless, the correlation between responses to platinum and PARP inhibitors is not absolute, emphasizing currently unappreciated distinctions in how DNA repair networks respond to these agents^79^.

The relative dispensability of ALC1 in cellular settings with intact DNA repair offers potential advantages over combinatorial therapeutic approaches that include cytotoxic agents with PARPi. ALC1 loss overcame several forms of engineered PARPi resistance and significantly sensitized BRCA1-mutant breast cancer xenografts that are modestly sensitive to PARPi. The cooperative activities of ALC1 and PARP on chromatin may also be relevant to a broad range of genomic damage, as demonstrated by the synergistic increase in IR sensitivity upon combining sg*ALC1* with PARPi (Fig. 7d-h). Notably, ALC1 loss by itself did not affect viability after IR, highlighting the expanded scope of DNA damage sensitivity when it is combined with low dose PARPi. These properties can potentially be exploited to overcome therapeutic resistance to radiotherapy and other clinically used modalities. Both the ATPase and macrodomains were essential for mediating PARPi response, thus presenting two moieties to develop small molecule inhibitors against. This approach of targeting chromatin accessibility offers new possibilities that extend beyond inactivation of a singular repair pathway. This enables an increased therapeutic window in response to PARPi and overcomes several forms of resistance that arise due to rewiring of DNA repair networks.

## Methods

### Cell culture

UWB1.289 and UWB1.289+*BRCA1* cells were purchased from ATCC and were maintained in 1:1 MEBM (Lonza): RPMI 1640 with GlutaMAX (Thermo Fisher) and supplemented with 10% fetal bovine serum (FBS, Atlanta Biologicals) and 1x penicillin and streptomycin (P/S, 100 U/ml, Gibco). DLD1 WT and DLD1 *BRCA2-/-* were purchased from Horizon Discovery and cultured in RPMI 1640 with GlutaMAX media and supplemented with 10% FBS and 1x penicillin-streptomycin (P/S). 293T and U-2 OS cells were purchased from ATCC and were maintained in DMEM media (Thermo Fisher) with 10% Bovine Calf Serum (BCS, GE HyClone) and 1x P/S. CAPAN1 cells were grown in DMEM with 10% FBS and 1x P/S. SUM149PT were cultured in Ham’s F-12 (Thermo Fisher) media supplemented with 5% FBS, 1x P/S, hydrocortisone (1 mg/ml, Sigma) and insulin (5 mg/ml, Sigma). hTERT-RPE1 *p53-/-* Cas9 and hTERT-RPE1 *p53-/-BRCA-/-* Cas9 cells were gifted by D. Durocher and were cultured in DMEM media with 10% FBS and 1x P/S. hTERT-RPE1 *p53-/-* Cas9, hTERT-RPE1 *p53-/- BRCA1-/-* Cas9, DLD1 WT and DLD1 *BRCA2-/-* cells were maintained at 3% oxygen, while other lines were maintained at atmospheric oxygen.

### Chemicals used in this study include

Olaparib (LC laboratories, 0-9201), talazoparib (Selleck Chemicals, S7048), camptothecin (Cayman Chemicals, 11694), H_2_O_2_ (Sigma, H1009), cisplatin (Tocris, 2251), MMS (Sigma, 129925), and PARGi (PDD 00017273, Fisher 5952). Most solid compounds were reconstituted in DMSO at 1000X of the required concentration, such that the final percentage of DMSO was 0.1%. To all untreated controls, a final concentration of 0.1% DMSO was added. Cisplatin was dissolved in water to yield a stock concentration of 5 mM.

### Vector construction and sgRNA cloning

Single knockouts were generated either using a two-vector *Streptococcus pyogenes* (Sp) Cas9 system: LentiV_SpCas9_puro (Addgene, 108100) and LRG2.1 backbone (Addgene, 108098) or an all-in-one (expressing both SaCas9 and sgRNA) single vector *Staphylococcus aureus* (Sa) Cas9 system. The SaCas9 all-in-one vector, henceforth referred to as SaLgCN, was derived by cloning the SaCas9 coding sequence (dCAS-VP64_Blast, Addgene: 61425) and its associated sgRNA expression cassette into a lentiviral vector. For double knockout experiments, ALC1 was knocked out using the single vector SaCas9 system and the two-vector SpCas9 system was employed for the depletion of the second protein of interest. Plasmids generated in this study, which will be available through Addgene: (1) SpCas9 sgRNAs expressed using LRG2.1 (Addgene, 108098), (2) modified version of LRG2.1 engineered to replace GFP with <u>b</u>lasticidin (LRG2.1<u>b</u>: U6-sgRNA-EFS-blast), (3) <u>n</u>eomycin (LRG2.1<u>n</u>: U6-sgRNA-EFS-Neo), and (4) SaLgC<u>N</u> (U6-sgRNA-EFS-SaCas9-P2A-<u>Neo</u>). sgRNAs were designed to target the functional domain of protein and were cloned by annealing the two complementary DNA oligos into a BsmB1-digested LRG2.1, LRG2.1b, LRG2.1n, or SaLgCN vector using T4 DNA ligase. To improve U6 promoter transcription efficiency, an additional 5’ G nucleotide was added to all sgRNA oligos that did not already start with a 5’ G. List of sgRNAs used in the study are listed in Supplementary Table 4.

### Lentivirus generation and transduction

Lentivirus were generated in HEK293T cells by transfecting VSVG, psPAX2 and the plasmid of interest in a molar ratio of 1:1:1.3 using polyethylenimine (PEI). Three mg of PEI was used for every mg of plasmid transfected. Media was changed 6-7 hours after transfection and virus was collected at 24, 48 and 72hrs post transfection and pooled. Virus was transduced on target cell lines by spin infection at 500 g for 30 min in the presence of polybrene (8 mg/ml). 16 hours post-transduction, media was changed and if required, antibiotic selection was performed 24 hours after virus removal. Puromycin (2 mg/ml), blasticidin (5 mg/ml), hygromycin (100 µg/ml) and neomycin (400 mg/ml) were used for selecting all cell lines except hTERT-RPE1, which were selected using puromycin (10 mg/ml) and neomycin (800 mg/ml).

For all biological replicates, independent lentivirus infection and antibiotic selection was performed each time.

### Competition-based cellular proliferation assays

Competition-based cellular proliferation assays were performed as previously described^80^. For the purposes of these growth competition experiments and CRISPR screen, T_initial_ was determined for each cell line. For a given cell line, T_inital_ denotes the day when the highest GFP expression was achieved for a sgRNA targeting an essential gene, for example, PCNA or RPA3. To determine the T_initial_, cells were transduced with sgRNA targeting an essential gene and GFP positivity was recorded for six consecutive days. Peak GFP was reached at Day 3 for hTERT-RPE1 (*p53*-/-, *BRCA1*-/-), Day 4 for UWB1.289 and UWB1.289+*BRCA1*, Day 5 for SUM149PT, DLD1 *BRCA2*-/- and CAPAN-1 and Day 6 for DLD1 WT. sgRNAs were cloned into a vector that also expressed GFP as a marker of transduction. Lentivirus were generated in an array format as described above. Cells were transduced at an MOI of <0.5 to allow competition between the GFP-positive cells (that carried the sgRNA) and GFP-negative cells. On the day of T_initial_, the percentage of GFP positive cells was recorded and cells were split into media containing either DMSO or olaparib. After every two population doublings, the percentage of GFP was measured and cells were passaged 1:4. GFP readings were taken for 6-7 passages (12-14 population doublings). Two population doubling time of cell lines used in the study was determined to be two days for hTERT-RPE1 *p53-/-*, DLD1 *WT*, three days for hTERT-RPE1 *BRCA1-/-*, four days for UWB1.1289+*BRCA1* and DLD1 *BRCA2-/-*, and five days for UWB1.289, SUM149PT and CAPAN1.

### Retrovirus plasmids and transduction

ALC1 cDNA was cloned into a retroviral pOZ-N vector as well as in a pMSCV-puromycin vector using Gibson Assembly (NEB, E5510S) with an N-terminus FLAG-HA tag. FLAG-HA XRCC1 was cloned into the pMSCV-puromycin vector using Gibson Assembly. Site-directed mutagenesis was performed using Q5 Site-Directed Mutagenesis Kit (NEB, E0552S). Retrovirus was generated by transfection of Phoenix cells with 3:1 ratio of target plasmid: pCL ampho DNA using LipoD293 (Fisher, SL100668). Virus were collected at 48 hr and 72 hrs after transduction and target cells were infected and centrifuged at 2000 rpm for 30 min. Selection for pOZ constructs was done using IL-2-conjugated magnetic beads and pMSCV-puromycin expressing cells were selected using 1 mg/ml of puromycin.

### Construction of domain-focused sgRNA pooled library

A gene list of chromatin regulators in the human genome was manually curated based on the presence of chromatin regulatory protein domains^32^. The chromatin regulatory protein domain sequence information was retrieved from NCBI Conserved Domains Database. Approximately 6 sgRNAs were designed against individual protein domains (Supplementary Table 5). The design principle of sgRNA was based on previous reports and the sgRNAs with the predicted high off-target effect were excluded^68^. All of the sgRNAs oligos including positive and negative control sgRNAs were synthesized in a pooled format (Twist Bioscience) and then amplified by PCR. PCR amplified products were cloned into BsmB1-digested LRG2.1 (a lentiviral sgRNA expression vector, U6-sgRNA-EFS-GFP, Addgene: 108098) using the Gibson Assembly kit (NEB#E2611). To verify the identity and relative representation of sgRNAs in the pooled plasmids, a deep-sequencing analysis was performed on a MiSeq instrument (Illumina) and confirmed that 100% of the designed sgRNAs were cloned into the LRG2.1 vector and the abundance of >95% of individual sgRNA constructs was within 5-fold of the mean. This chromatin regulatory domain CRISPR sgRNA pooled library will be available through Addgene.

### CRISPR-based pooled library screening

CAPAN-1, SUM149PT, UWB1.289 and UWB1.289+*BRCA1* were engineered to stably express SpCas9 (a lentiviral expression vector, EFS-Cas9-P2A-Puro vector, Addgene: 108100). Lentivirus of pooled sgRNA library targeting functional domains of chromatin factors was generated as described above. To ensure a single copy sgRNA transduction per cell, the multiplicity of infection (MOI) was set to 0.3-0.4. At T_initial_ a fraction of cells were collected to prepare the reference representation and the rest were propagated either in the presence of DMSO or 10 nM olaparib, which approximately corresponds to the lethal dose 20 (in a two-week clonogenic assay) of the cells used for screening. Cells were passaged for 14 population doublings and were harvested at the final time point. To maintain the representation of sgRNAs during the screen, the number of sgRNA positive cells was kept at least 1000 times the sgRNA number in the library. Genomic DNA was isolated using QIAamp DNA Mini and Blood Mini kit (QIAGEN). sgRNA cassettes were PCR amplified from the genomic DNA using Phusion Flash High Fidelity PCR Master Mix (Thermo Scientific, catalog#F548L). The PCR product was then end-repaired to produce blunt ends using T4 DNA polymerase (NEB M0203L), DNA Polymerase I, Large (Klenow) Fragment (NEB M0210L), and T4 polynucleotide kinase (NEB M0201L). 3’ A-overhang was then added to the ends of blunted DNA fragments with Klenow Fragment (3’-5’ exo-) (NEB M0212L) and the PCR clean-up was performed using QiaQuick kit. The DNA fragments were ligated to Bar Code Adapters using Quick Ligation kit (NEB M2200L) and purified using Agentcourt AMPure Beads (Beckman Coulter, A63880). Next, Illumina paired-end sequencing adaptors were attached to the barcoded ligated products through PCR reaction with Phusion Flash High Fidelity Master Mix (Thermo Fisher, catalog#F531S). The purified product was quantified by Bioanalyzer Agilent DNA 1000 (Agilent 5067-1504) and pooled together in equal molar ratio and pair-end sequenced by using MiSeq (Illumina) with MiSeq Reagent Kit V3 (150 cycle) (Illumina, MS-1-2-3001). The sequencing data was de-multiplexed and trimmed to contain only the sgRNA sequence cassettes. The read count of each individual sgRNA was calculated with no mismatches and compared to the sequence of reference sgRNA as described previously^25^. Individual sgRNAs with the read count lower than 50 in the initial time point were discarded and remaining sgRNA counts normalized to total sample read counts. A protein domain CRISPR Score (CS) was calculated by averaging the log2 fold-change of all CRISPR RNA targeting a given protein domain. Log2 fold-change=(final CRISPR RNA abundance + 1)/(initial CRISPR RNA abundance). The log2 fold-change values for each protein domain are provided in Supplementary Table 1.

### Clonogenic assay

For clonogenic experiments with SUM149PT, UWB1.289 and UWB1.289+*BRCA1*, 500 cells were plated in technical triplicates and analyzed after 10-14 days. For hTERT-RPE-1, 250 cells were plated in a 10 cm dish. hTERT-RPE1 *p53-/-* type and p53-/- *BRCA1-/-* cells were grown at 3% oxygen for 8 and 11 days, respectively. For DLD1 cells, 500 cells were plated in triplicates in 6-well cm plates. Both WT and BRCA-mutant DLD1 cells were grown at 3% oxygen for 9 and 14 days respectively. For cellular complementation analysis with mutants, cells were directly plated in media containing olaparib (0.5 or 1 nM). For clonogenic experiments with PARGi, cells were plated on Day 0, PARGi was added on Day 1 and olaparib was added after a 24 hr incubation with PARGi. For IR+PARPi experiments, cells were plated on Day 0, treated with PARPi on Day 1 and IR treatment was performed 24 hrs after PARPi addition. After 14 days cells were washed with PBS and stained with 0.4% crystal violet in 20% methanol for 30 min at room temperature. Plates were then washed with deionized water and air-dried. Colonies were manually counted. Bliss scores for analyzing combinatorial responses were generated using Combenefit^81^.

### CellTiter-Glo assay

1000 cells in a volume of 100 µl were plated into each well of a 96-well clear bottom black plates (Corning, Neta Scientific 3904) on Day 0. On the next day, 2X drug dilutions were made and 100 ml of the drug was added to cells in technical triplicates. While other drugs were retained, MMS-treated cells were released into fresh media after 24 hrs. The viability was measured using CellTiter-Glo Luminescent Cell Viability Assay (Fisher, G7572) either after 5 days of drug addition (for RPE-1 and U-2 OS) or after 7 days (for DLD1, UWB1.289, UWB1.289+BRCA1 and SUM149PT). When plotting the survival curves, the luminescence of the drug-treated population was normalized to solvent-exposed cells. The data was fitted in GraphPad Prism using the following equation: y = min+ (max-min)/1+10^logEC50-x^.

### Xenograft experiment

Xenograft studies were carried out under protocol number 803170 approved by Institutional Animal Care & Use Committee at the University of Pennsylvania. 5-week-old female NSG (NOD.Cg-*PrkdcscidIl2rgtm1Wjl*/SzJ) mice were procured from Jackson Laboratory and tumor implantation was performed once these mice were 7-weeks old. SUM149PT-Cas9 cells were engineered to stably express either sgNeg or sgALC1. 5 days after transduction (T_initial_ for SUM149PT), 2 x 10^6^ cells mixed 1:1 with Matrigel (Fisher Scientific, 356237) and were subcutaneously implanted into both flanks of the mice. Once the tumor volume reached between 80-130 mm^3^, mice were randomized into vehicle and drug treated groups, such that the average tumor volume was approximately 100 mm^3^. Olaparib (LC laboratory) was dissolved in DMSO to a concentration of 50 mg/ml and was further diluted 1:10 in 15% hydroxypropyl b-cyclodextrin (Sigma) (w/v in PBS 7.4). 50 mg/kg of the drug was administered for five consecutive days of a week using oral gavage. 1:10 dilution of DMSO in 15% hydroxypropyl b-cyclodextrin (w/v in PBS 7.4) was used as vehicle control. Tumor dimensions were measured twice a week using a digital caliper and the volume was calculated using the formula: (length × width^2^)/2. Mice were euthanized once the tumor size reached >10.5 mm in any direction. The harvested tumors were snap frozen in liquid nitrogen for western studies.

### Metaphase spreads

Cells growing at 70-80% confluency were treated with colcemid (100 ng/ml) for 3 hrs. Cells were harvested by trypsinization. The cell pellet was uniformly suspended in KCl (0.075 M, pre-warmed at 37°C) followed by incubation at 37°C for 30 min with intermittent mixing. Cells were collected by centrifugation and fixed by dropwise addition of ice-cold methanol: acetic acid (3:1) and incubated overnight. Methanol pre-soaked slides were kept on a humidified heat block at 42°C and cells were dropped from a height and were spread by air blowing. Slides were stained with Giemsa for 3-6 minutes, washed, and mounted using Permount.

### Immunofluorescence

PARP2 trapping was performed using the methodology described by Michelena et al. ^45^. Briefly, cells were grown in 8-well chamber slides (Fisher, 154941PK). Post-treatment, cells were pre-extracted on ice for 5 min using Triton X-100 (0.2%) in PBS. Cells were then fixed with 4% paraformaldehyde (PFA) for 15 min at RT, followed by PBS washes. Cells were then permeabilized at room temperature for 5 min using Triton X-100 (0.2%) in PBS. DMEM/FBS was used for blocking and making antibody dilutions. After blocking cells for an hour at RT, cells were incubated with 1:200 dilution of PARP2 antibody (Active Motif, 39743) overnight at 4°C. Following washes with Tween (0.2 %) in PBS (PBST), cells were incubated with 1:200 anti-rabbit secondary for an hour at room temperature, followed by 3-4 washes with PBST and mounting in Vectashield Antifade Mounting Medium with DAPI (Vector Laboratories, H-1200). Images from two biologically independent experiments were captured at the same time using Zeiss Axio Widefield (20x/0.8) microscope. Immunostaining results were analyzed using CellProfiler 3.0 software, whereby a specialized pipeline was implemented to identify and measure PARP-2 and DAPI signal intensity (*CellProfiler Module: IdentifyPrimaryObjects/MeasureObjectIntensity*). Illumination correction was applied to each image prior to intensity measurements to account for non-uniformities in illumination. In addition, PARP-2 signal was masked with DAPI-stained nuclei (*CellProfiler Module:MaskImage – nuclei object*) to ensure that only nuclear PARP-2 signal intensity was measured. From this analysis, an output measure of *IntegratedIntensity*/per object/per field-of-view was obtained for each experimental condition and plotted using GraphPad Prism software.

To examine γH2AX foci during S-phase, cells were grown on poly-L-lysine (Poly-K) (Sigma, P4832) coated coverslips and were treated with 10 mM EdU for 20 min before harvesting. Cells were pre-extracted on ice for 5 min using Triton X-100 (0.5%) in CSK buffer (10 mM PIPES pH6.8, 100 mM NaCl, 300 mM Sucrose, 3 mM MgCl2 and 1 mM EGTA) and fixed using PFA (4%) in PBS for 15 min at room temperature (RT). Following 3-4 washes with PBS, cells were permeabilized using Triton X-100 (0.5%) in PBS (PBSTx) for 5 min on ice and blocked using BSA (3%) in PBS for 30 min at room temperature. Alexa Fluor dye was conjugated to EdU by click chemistry using Click-iT Cell Reaction Buffer kit (C10269, Thermo Fisher). After washes with PBST, cells were fixed again with PFA (4%) for 10 min. Following PBS washes, cells were incubated with a 1:2000 dilution of anti-phospho-Histone H2A.X (Ser139) clone JBW301 antibody (Millipore, 05-636-1) for an hour at room temperature. Cells were washed with PBST and were next incubated with the Alexa Fluor conjugated mouse secondary (1:1000, Life Technologies) for an hour at RT. Coverslips were mounted using the Vectashield with DAPI. Images were acquired using a CoolSnap Myo camera (Photometrics) connected to the Nikon NIS-Elements software. Images were processed using Fiji (National Institutes of Health), numbers of γH2AX foci were determined using Find Maxima function and statistical analysis was performed using GraphPad Prism.

For quantifying RAD51-γH2AX co-localization, cells grown on Poly-K coverslips were pre-extracted on ice for 3 min using Triton X-100 (0.5%) in CSK buffer and fixed using PFA (4%) in PBS for 15 min at room temperature. Following washes with PBS and permeabilization with PBSTx, cells were blocked using blocking buffer (5% normal donkey serum, 0.1% fish skin gelatin, 0.1% Triton X-100, 0.05% Tween-20, and 0.05% sodium azide in PBS) for 1 hour at RT. Cells were incubated overnight with 1:200 dilution of RAD51 antibody (H-92, sc-8349, Santacruz) and 1:2000 dilution of anti-phospho-Histone H2A.X (Ser139) clone JBW301 antibody made in blocking buffer. Following washes with PBST, cells were incubated with 1:200 dilution of Alexa Fluor 488 conjugated anti-rabbit and 1:1000 dilution of Alexa Fluor 568 conjugated anti-mouse secondary (Life Technologies) for an hour at RT, mounted and imaged as described above. Co-localization events were manually counted keeping brightness and contrast constant throughout the analysis.

For quantifying ss-DNA, cells growing on Poly-K coverslips were labeled with CIdU (10 mM, C6891, Sigma) for 48 hours. After pre-extraction on ice for 5 min using Triton X-100 (0.5 %) in CSK buffer, cells were fixed in PBS containing PFA (4%) for 15 min at room temperature, washed with PBS, and blocked using blocking buffer for an hour at room temperature. Cells were incubated with 1:100 dilution of CIdU antibody (Abcam, ab6326) and 1:2000 dilution of Anti-Histone H2A.X (phosphor S139) [EP854(2)Y] (Abcam, ab81299) for an hour at 37°C. This was followed by washes with PBST and incubation with Alexa Fluor 488 conjugated anti-mouse (1:200 dilution) and Alexa Fluor 568 conjugated anti-rabbit secondary antibodies (1:1000 dilution) for an hour at RT. Coverslips were mounted and imaged as described above. CIdU foci that did not co-localize with γH2AX were manually counted, keeping the values for image brightness and contrast constant throughout the analysis.

To quantify ALC1 and XRCC1 on chromatin, cell lines stably expressing HA-ALC1 or HA-XRCC1 were grown on Poly-K coverslips. After indicated treatment, cells were pre-extracted on ice for 2 min using Triton X-100 (0.5%) in CSK buffer and fixed using PFA (4%) in PBS for 15 min at room temperature. Coverslips were then washed with PBS and permeabilization using ice cold Methanol: Acetic Acid (1:1) on ice. Cells were blocked using DMEM/FBS media for 1 hour at RT followed by incubation with 1:1000 dilution mouse anti-HA (Biolegend, 901514) and rabbit anti-PCNA (Cell Signaling, 13110S) for 1 hour at RT. Following washes with PBST, cells were incubated with 1:200 dilution of Alexa Fluor 488 conjugated anti-mouse and 1:200 dilution of Alexa Fluor 568 conjugated anti-rabbit secondary (Life Technologies) for an hour at RT, followed by washes and mounting in Vectashield with DAPI. To quantify the chromatin bound signal, the DAPI channel was used to define the region of interest (ROI) using the magic wand tool in Image J. Integrated density for the defined ROI was then computed using the command M function.

### S1 nuclease fiber labeling

S1 nuclease fiber labeling was performed as described before^82^. 0.125 x 10^6^ UWB1.289 cells were plated in a 6-well plate on Day 0. On day 1, olaparib (100 nM) treatment was initiated for 24 hrs. 24 hrs post treatment, cells were pulsed with CIdU (25 mM, C6891, Sigma) for 20 min followed by a PBS wash and then pulsed for 20 min with IdU (250 mM, I7125, Sigma). Cells were then washed with PBS and permeabilized using CSK buffer (100 mM NaCl, 10 mM MOPS pH 7, 3mM MgCl_2_, 300 mM Sucrose and 0.5% Triton X-100) for 10 min at RT. After a gentle wash with PBS, digestion with S1 nuclease (20 U/ml) in S1 nuclease buffer (30 mM sodium acetate, 10 mM zinc acetate, 5% glycerol and 50 mM NaCl) was performed for 30 min at 37°C. Cells were then collected in PBS+0.1% BSA by scraping, spun at 2000 rcf for 7 min to harvest the cell pellet and then re-suspended in 50-100 ml of PBS+0.1% BSA. 3 ml of cells suspension was mixed with 12.5 ml of spreading buffer (200 mM Tris pH7.4, 50 mM EDTA, 0.5% SDS) on positively charged slides (Stellar Scientific, 1358B), incubated for 10 min followed by spreading the DNA on slides by titling them at a 45° angle. Samples were fixed using methanol: acetic acid (3:1) overnight. On the next day, slides were rehydrated and DNA was denatured using HCl (2.5 N) for an hour followed by PBS washes until the pH reached 7.5. Blocking was then performed with BSA (3%) in 0.1% Tween-80 in PBS (0.1% PBST) for an hour. CIdU and IdU were detected using mouse anti-IdU (BD-347580) and rat anti-CIdU (Serotec-OBT0030G) antibodies. After washes with 0.1% PBST, slides were incubated with Alexa Fluor conjugated anti-mouse and anti-rat antibodies (1:200 dilution) and mounted using the Vectashield without DAPI (Vector Laboratories, H-1000). Images were acquired as described above. Only fibers with both red and green signals were analyzed. The line-scan function of ImageJ was used to calculate the length of the fibers.

### Chromatin fractionation

Chromatin fractionation was performed using the Thermo Scientific Subcellular Protein Fractionation Kit for Cultured Cells (Fisher, 78840) following the manufacturer’s instruction. Modifications in the protocol included scraping cells in cold PBS supplemented with PARPi used for the respective experiment and MNase digestion was performed at 37°C for 15 min. An extra wash step with nuclear extraction buffer was introduced before MNase digestion.

### Immunoprecipitation

Exponentially growing HEK293T cells were transfected in a 10 cm dish with 3x FLAG-tagged ALC1 constructs using 1:3 ratio of DNA:PEI. 48 hours after transfection, cells were scrapped in cold PBS and the cell pellet was collected after spinning down at 500g for 5 minutes at 4°C. Cell pellets were suspended in lysis buffer (1 mL, 50 mM Tris, pH8, 137 mM NaCl, 1% Triton X-100, 1 mM MgCl_2_, 1 mM EDTA) supplemented with 1x protease inhibitor cocktail (Roche), PMSF (1 mM, Sigma), DTT (1 mM, Sigma) and benzonase (50 units/mL, 70746-3, Sigma) and incubated on an end-to-end rotor for 1 hour at 4°C. Cell lysates were then collected after centrifugation at 20,000 g for 15 min at 4°C. 15ul of lysis buffer equilibrated FLAG-M2 beads (A2220, Sigma) was incubated with 2mg of lysates for another hour at 4°C on an end-to-end rotor. The beads were then washed 4x times using the protease inhibitor and DDT supplemented lysis buffer. Bound proteins were eluted by adding 2X NativePAGE^TM^ sample buffer (BN2003) and heating at 95°C for 10 min. before analyzing by immunoblotting as described below.

### Immunoblotting

Cells were lysed using 2-5 times the volume of RIPA lysis buffer supplemented with 1x protease inhibitor cocktail (Roche), PMSF (1 mM, Sigma), DTT (1 mM, Sigma). The supernatant was collected after centrifugation and quantified using Bradford (Bio-Rad). Frozen tumors harvested from mice were minced and submerged into 2-5 volumes of RIPA buffer supplemented as above. Samples were homogenized by sonication on ice and incubated for 30 min at 4°C while shaking. Supernatant was then collected by centrifugation and quantified using Bradford. 15-20 mg of total cell, tumor or chromatin lysates were analyzed on NuPAGE 4-12% Bis-Tris gels (Invitrogen) and transferred to a nitrocellulose membrane (GE, 10600004). All western blots were performed using the Mini Trans-Blot apparatus (Bio-rad, 1703930) at 400 mA for 2 hours on ice. Blots were blocked in 5% milk in PBST and then incubated overnight with primary antibody at 4°C. After washes with PBST, blots were incubated with 1:3000 dilution of ECL anti-mouse or rabbit HRP linked secondary at RT for 1 hour, washed with PBST and developed using Western Lightning Plus-ECL (PerkinElmer, NEL105001EA) on autoradiography films. All unprocessed scans of blots are provided in Supplementary Fig. 1.

### Antibodies used for immunoblotting

ALC1 (Santacruz Biotechnology, sc-81065, 1:200 dilution), XRCC1 (Abcam, ab1838, 1:200 dilution), PARG (Thermo Fisher Scientific, PA5-14158), CHD4 (Proteintech, ab264417), APE-1 (Santacruz Biotechnology, sc-17774, 1:200 dilution), NTHL1 (Santacruz Biotechnology, sc-2660C1a, 1:200 dilution), 53BP1 (Novus Biological NB100-94, 1:1000 dilution), PARP1 (Cell Signaling mAB#9532, 1:2000 dilution), PARP2 antibody (Active Motif, 39743, 1:1000 dilution), HA (Biolegend, 901514, 1:1000 dilution), GFP (Cell Signaling mAb#2956, 1:1000 dilution), Rev7 (Invitrogen PA5-49352, 1:1000 dilution), GAPDH (Cell Signaling 2118S, 1:2000 dilution), mouse Anti-FLAG (F1804, Sigma), anti-Histone H4 (Millipore, 05-858).

### Assay for ATAC-seq

ATAC-seq was performed as previously described^83^. Reads from ATAC-seq experiments were trimmed with Trim Galore (version 0.4.1) with parameters -q 15 --phred33 --gzip --stringency 5 -e 0.1 --length 20. Trimmed reads were aligned to the Ensembl GRCh37.75 primary assembly including chromosome 1-22, chrX, chrY, chrM and contigs using BWA (version 0.7.13) (Li and Durbin, 2009, DOI: 10.1093/bioinformatics/btp324) with parameters bwa aln -q 5 -l 32 -k 2 -t 12 and paired-end reads were group with bwa sample -P -o 1000000. Reads mapped to contigs, ENCODE blacklist and marked as duplicates by Picard (version 2.1.0) were discarded and the remaining reads were used in downstream analyses. Peaks in each sample were identified using MACS (version 2.0.9) with parameters -p 1E-5 --nomodel --nolambda --format=BAM -g hs --bw=300 --keep-dup=1. A union of all peaks in untreated wildtype, PARPi treated wildtype, untreated ALC1 KO and PARPi treated ALC1 KO samples were generated using bedtools (version 2.27.1) ‘merge’ function.

To determine the change in accessibility, aligned reads of each ATAC-seq sample were quantified on the union of peaks and normalized to FPKM. Log2 fold change of accessibility was calculated as log2 averaged FPKM on replicates of other conditions versus untreated wildtype. Significance of change was determined using unnormalized quantification as input to ‘DESeq’ function from DESeq2 package in R (version 3.6.1) with parameters test = “Wald”, betaPrior = F, fitType = “parametric”. *P*-values were adjusted for multiple hypothesis testing using FDR. The average ATAC-seq signal +/− 1Kb round the center of the union of peaks were calculated using ‘metagene’ R package and metagene plot was generated using ‘ggplot2’ R package. One-tailed paired t-test of the average profiles was performed with other conditions versus untreated wildtype using ‘t.test’ function in R with paramters paired = ‘TRUE’ and alternative = ‘less’.

### RNA-seq

BRCA1-mutant hTERT-RPE1 cells and BRCA2-mutant DLD1 cells were infected with sgRNAs targeting *ALC1* in a lentiviral format and 1-5 million cells were harvested at the 4^th^ population doubling time after their respective T_inital_. Cells were lysed in TRIzol (Invitrogen) and RNA was isolated using Direct-zolTM RNA Miniprep Plus kit (ZYMO). RNA quality was assessed by RNA 600 Nano Bioanalyzer kit (Agilent) and samples with RIN ≥ 9.5 were used for library construction. RNA-seq libraries were prepared with QuantSeq 3′mRNA-Seq Library Prep Kit FWD for Illumina (Lexogen) by using 2 µg of total RNA as input and 11 cycles for PCR amplification. Library quality was checked by High Sensitivity DNA Bioanalyzer kit (Agilent) and libraries were sequenced on NextSeq 500/550 Platform with 75 bp single-end reads.

### RNA-seq analysis

Raw reads were mapped to the human (hg38) genome using Lexogen Quantseq 2.3.1 FWD platform with low quality reads removed. Raw reads were mapped into the hg38 genome using STAR Aligner with modified ENCODE settings. Gene raw read count files were generated with HTSeq-count. Differentially expressed genes were identified using DESeq2 (1.14.1) with default settings, and raw read counts >2 in all conditions were considered expressed. Genes with |log2FC| ≥ 0.5 and an adjusted *p* value <0.05 were considered significant and strongly altered.

### Statistics

All statistical analyses were performed using GraphPad Prism 8 software. Significance was calculated either by the two-tailed student’s *t*-test or Mann-Whitney test unless otherwise specified.

### Data availability statement

Sequencing data generated in this study has be deposited in the Gene Expression Omnibus with accession code # GSE149104 (for RNA-seq) and # GSE150955 (for ATAC-seq). Data from the CRISPR screen has been provided as mapped reads in Supplementary Table 1. Numerical source data and replicates have been provided in Supplementary Table 6.All other source data from this study are available from the corresponding author upon reasonable request.

### Code availability statement

All the analyses were based on standard algorithms described in the methods and referenced accordingly. There are no custom algorithms to make available.

## Acknowledgements

We thank D. Durocher (Univ. Toronto, Lunenfeld) for sharing hTERT-RPE1 *p53-/-BRCA1-/-* cells and controls, J. and R. Connaway (Stowers) for ALC1 plasmids, Keith Caldecott (Univ. of Sussex) for sharing GFP-XRCC1 plasmid and hTERT-RPE1 *XRCC1-/-* and parental control cells, Nick Lakin (Univ. of Oxford) for sharing *U-2 OS PARP1-/-*, *PARP2-/-* and *PARP1/2-/-* cells, and N. Johnson (Fox Chase) for SUM149PT BRCA1 reversion mutant cell lines and helpful discussions on mouse xenograft experiments. This work was supported by NIH grants GM101149 and CA17494 (to RAG), who is also supported by funds from the Penn Center for Genome Integrity, the Basser Center for BRCA, a V Foundation Team Convergence Award, and a Gray Foundation Team Science Award. PV was supported by the Ann and Sol Schreiber Mentored Investigator Award (Ovarian Cancer Research Fund Alliance) and Pilot funds from the Ovarian Cancer Translational Center for Excellence (UPenn). JS was supported by the Michele and Kevah Konner Award through the Basser Center for BRCA.

## Authors Contributions

P.V., J.S. and R.A.G. designed the study; P.V. did most of the experiments, with assistance from P.V.D, M.D., Y.S., Y.L. and S.P; Z.C and E.A. performed the CRISPR screen in SUM149PT and CAPAN1 cells and the RNA seq experiments; Y.Z. did the ATAC-seq experiment under the guidance of R.B.F.; W.L. performed the mouse work; L.P. imaged and analyzed PARP2 trapping under the guidance of R.H.M., P.V. and R.A.G. wrote the manuscript with contributions from J.S. R.A.G. is a founder and scientific advisory board member of RADD Pharmaceuticals.

---

**Extended Data Fig. 1.**
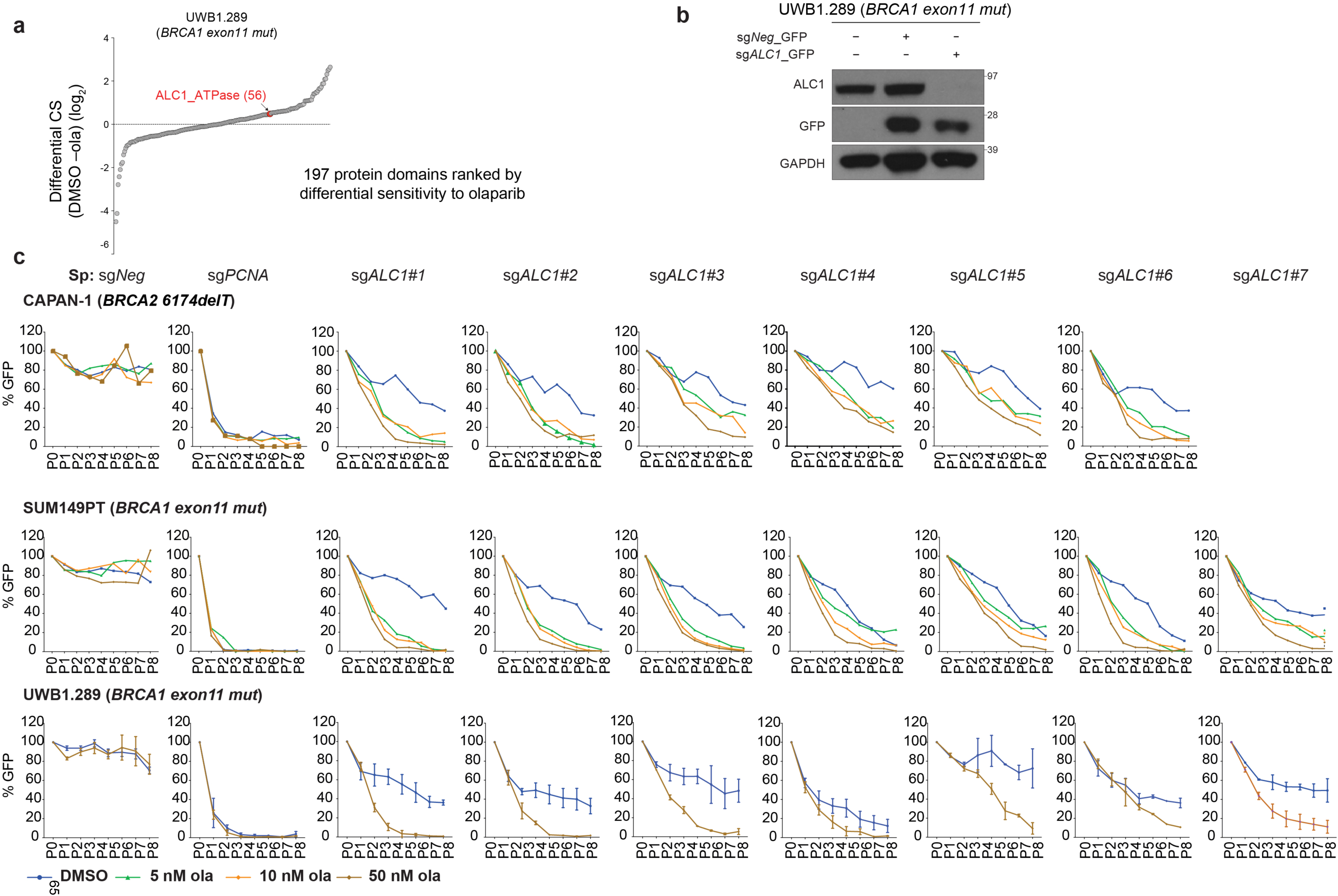
ALC1 loss renders olaparib hypersensitivity and proliferation defects in various BRCA-mutant lines. **a**, Protein domains ranked on the basis of the CRISPR score for ola sensitivity in BRCA1 mutant UWB1.289 cells. **b,** Immunoblot examining depletion of ALC1 using sgRNA vector with a GFP selection marker. GFP positive cells were sorted and analyzed. **c**, GFP competition assay to examine the effects of ALC1 depletion on ola sensitivity in CAPAN-1, SUM149PT and UWB1.289 cells using different sg*ALC1* guides.

**Extended Data Fig. 2.**
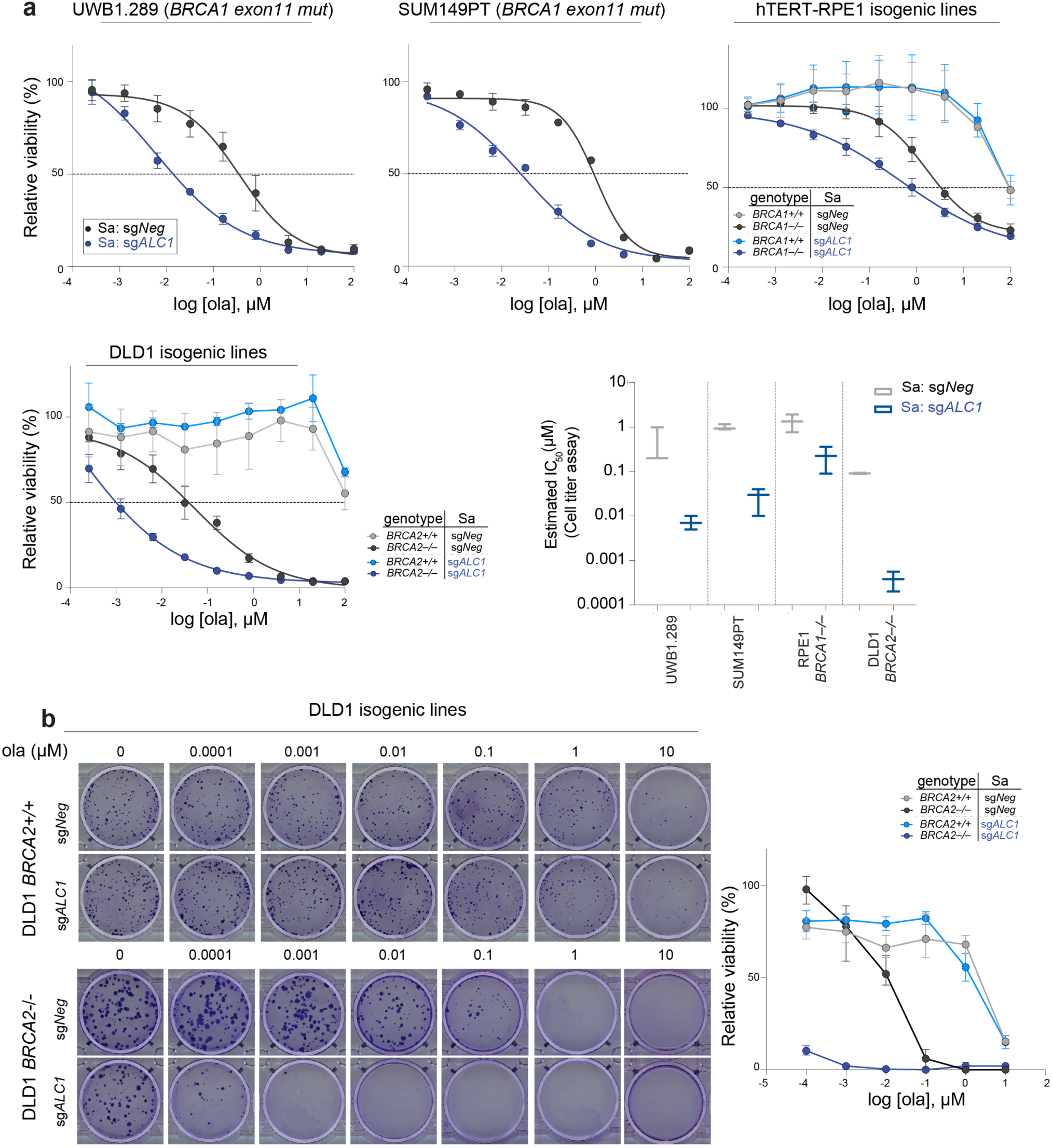
ALC1 loss enhances the therapeutic window of olaparib sensitivity in BRCA-mutant cells. **a**, Sensitivities of the indicated cell lines to ola using CellTiter-Glo. Data are mean ± s.e.m. from 2 (DLD1 and hTERT-RPE1) or 3 (UWB1.289 and SUM149PT) biologically independent experiments. **b,** Representative images (left) and quantification (right) of the clonogenic survival of ALC1-depleted DLD1 WT and BRCA2 mutant cells grown in the presence of increasing concentrations of ola.

**Extended Data Fig. 3.**
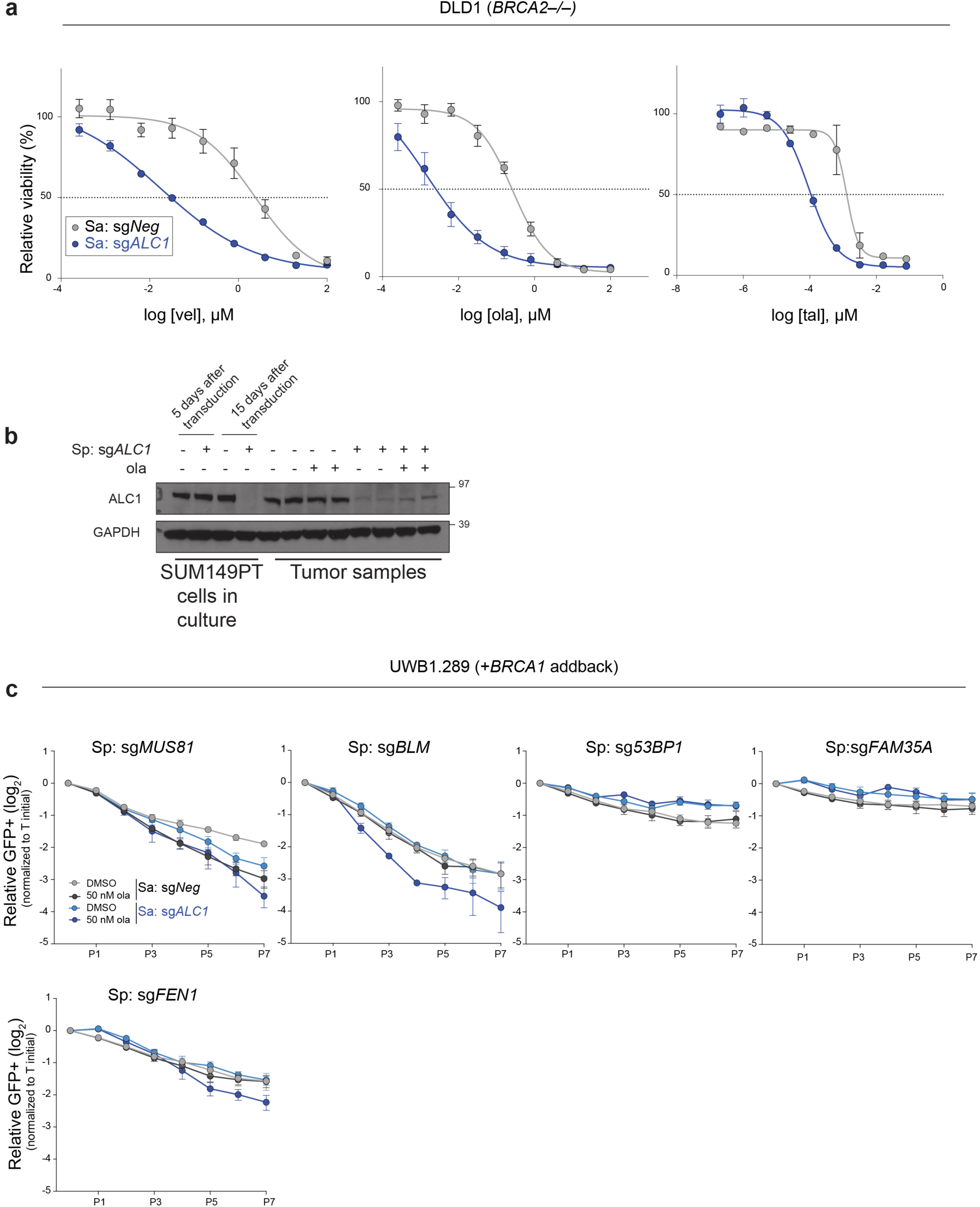
Extended analysis of PARPi sensitivity upon ALC1 loss. **a**, Sensitivities of the indicated DLD1 BRCA2-/- cells to vel (veliparib), ola and tal in CellTiter-Glo assay. Data are mean ± s.e.m. from 2 (tal) and 3 (vel and ola) biologically independent experiments. **b**, Immunoblot showing ALC1 levels in cells used for xenograft studies (first four left lanes) and in tumors that reached >10.5 mm in any dimension, which is when the mice were euthanized. **c,** GFP competition experiment in UWB1.289+*BRCA1* addback line to examine the effects of the combined loss of ALC1 and the indicated DNA repair proteins on cell proliferation and ola sensitivity. Data are normalized to Tinitial and indicate mean ± s.e.m. After every two population doublings, cells were passaged (P) and GFP percent was recorded (n=2-6 independent transductions).

**Extended Data Fig. 4.**
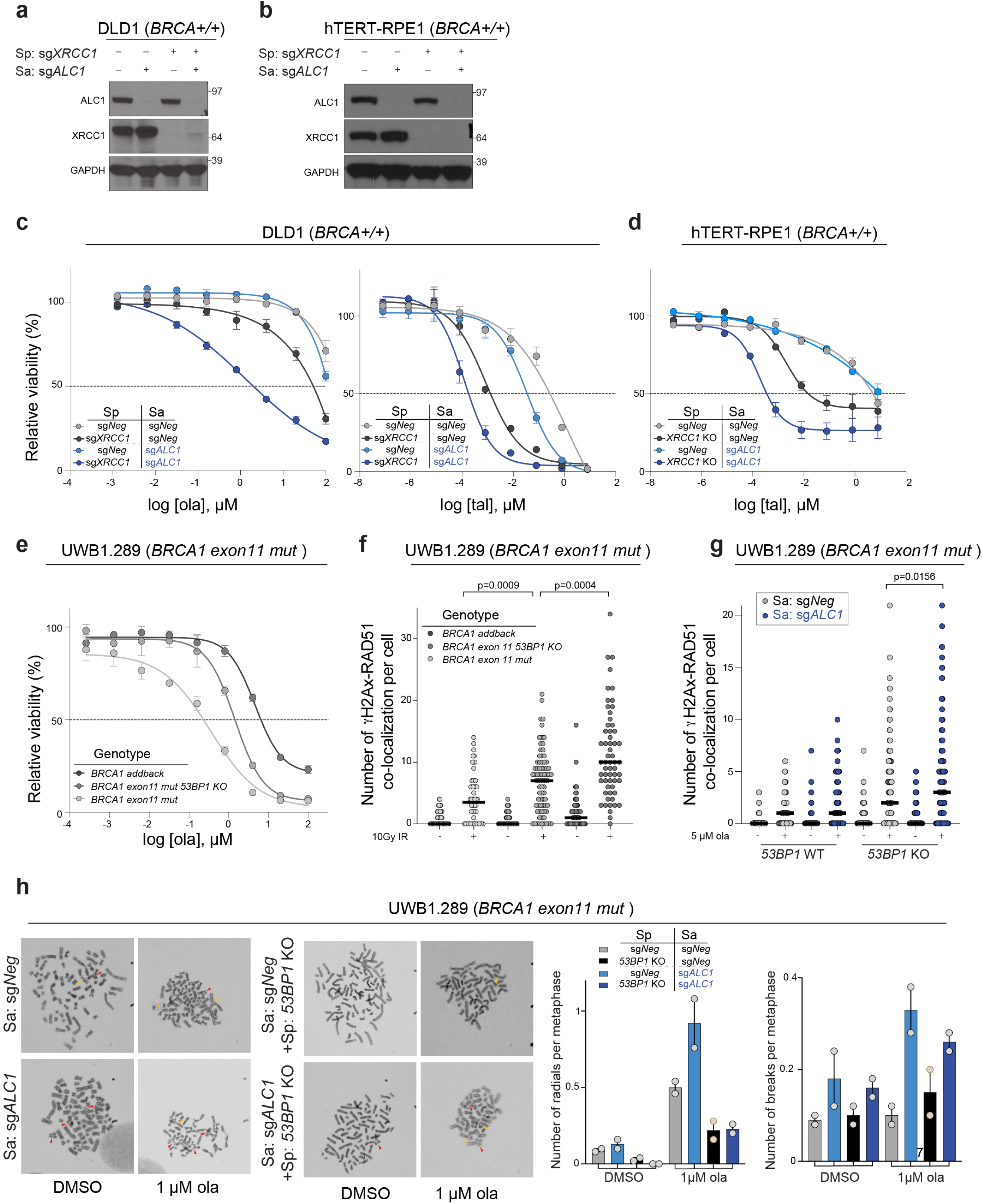
Genomic lesions in PARPi treated ALC1 deficient cells are repaired by SSBR and NHEJ. **a-b,** Immunoblot showing levels of ALC1 and XRCC1 in indicated DLD1 **(a)** and hTERT-RPE1 **(b)** cells **c,** Sensitivities of the indicated DLD1 cells lines to ola and tal using the CellTiter-Glo assay. Data are mean ± s.e.m. from three biologically independent experiments. **d,** Sensitivities of the indicated hTERT-RPE1 cells lines to tal using the CellTiter-Glo assay. Data are mean ± range from two biologically independent experiments. **e,** Sensitivities of the indicated UWB1.289 cells lines to ola using the CellTiter-Glo assay. Data are mean ± s.e.m. from three biologically independent experiments. **f,** Quantification of γH2AX-Rad51 foci in indicated cell lines. Cells were fixed 16 hrs after treatment with 10 Gy ionizing radiation (IR). Median is indicated. *P* value, Mann-Whitney. **g,** Quantification of γH2AX-Rad51 foci in indicated cell lines. Cells were treated with 5 µM ola for 24 hours before fixation. Data are from three biologically independent experiments. Median is indicated, *p* value, Mann-Whitney. **h,** Representative images and quantification of radials (indicated by red arrow heads) and breaks (indicated by yellow arrowheads) in indicated UWB1.289 cell line, post treatment with 1 µM ola for 24 hrs. For each experiment, at least 50 spreads were analyzed per sample. Data are mean ± range from two biologically independent experiments.

**Extended Data Fig. 5.**
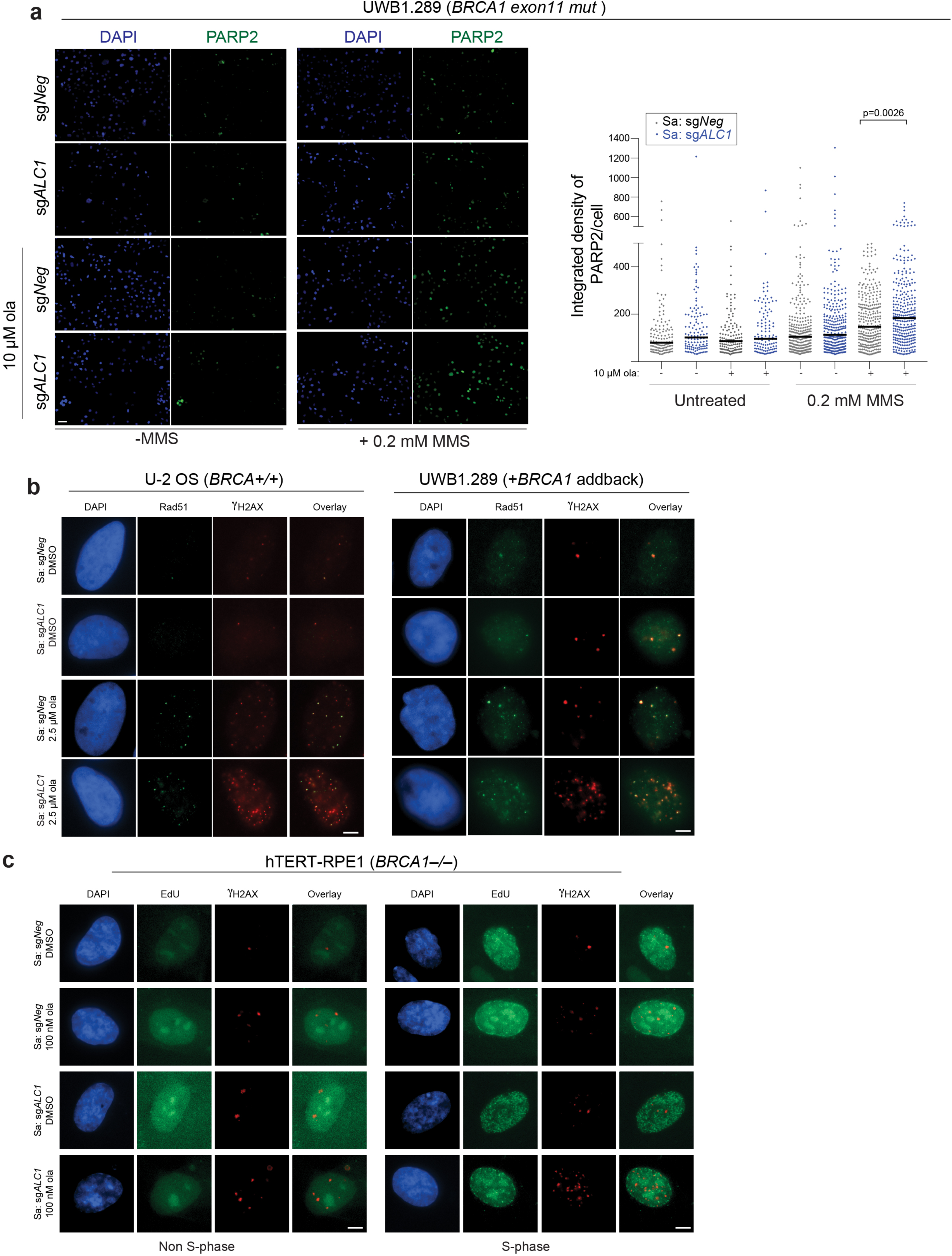
ALC1 deficiency results in increased PARP2 trapping and replication-coupled damage. **a**, Representative images (left) and quantification (right) of PARP2 trapping in UWB1.289 cells. Indicated treatments were performed for 4 hours. Data are from two biologically independent experiments. For each experiment, at least 50 cells were quantified per sample, *p* value, Mann-Whitney. Scale bar: 50 microns. **b,** Representative images of Rad51-γH2AX foci in U-2 OS (left) and UWB1.289+BRCA1 (right) cell lines. **c,** Representative images of γH2AX signal in Non S-phase (left) and S-phase (right) hTERT-RPE1 cell lines. Scale bar: 10 microns.

**Extended Data Fig. 6.**
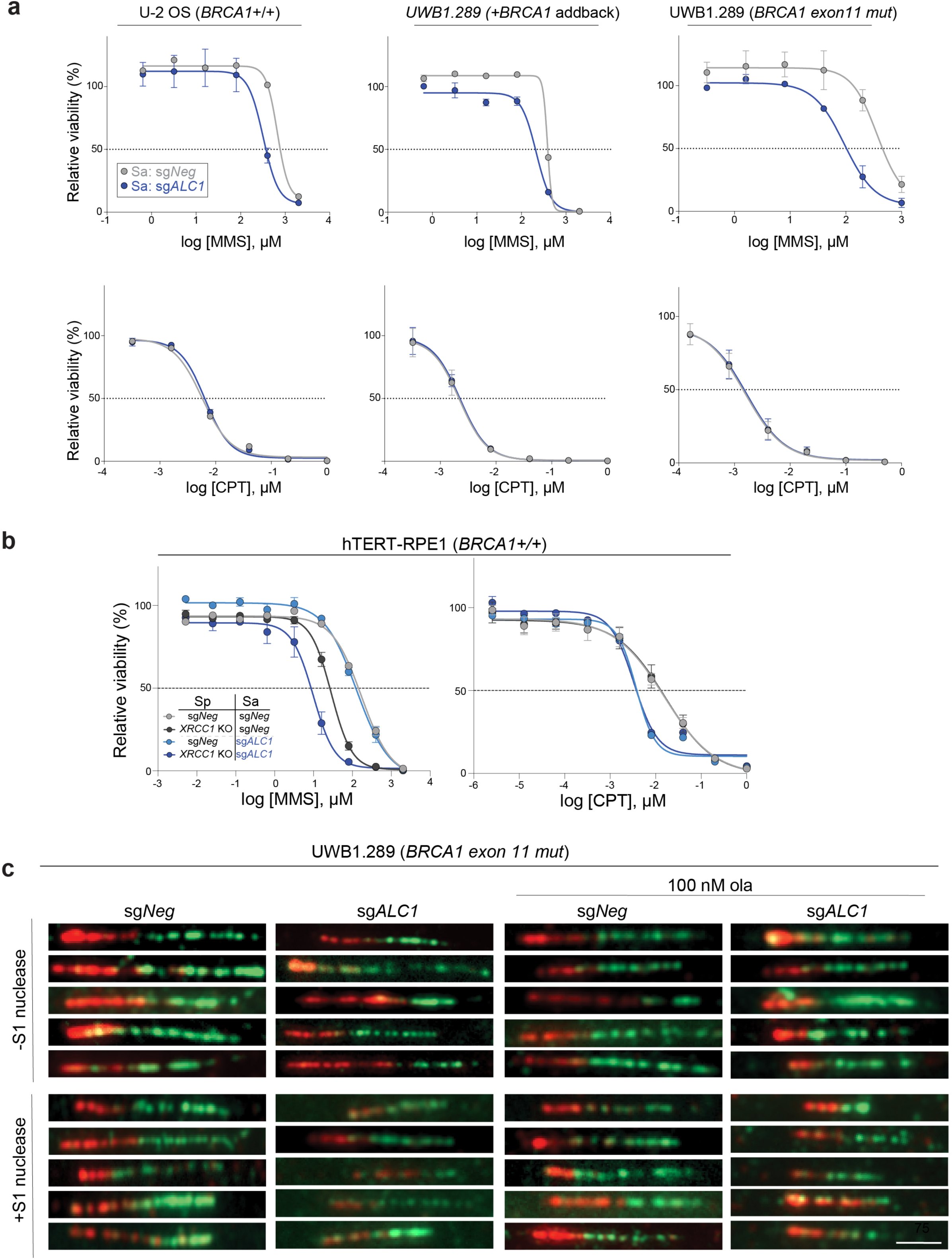
ALC1 loss confers MMS sensitivity and results in replication-coupled gaps. **a,** Sensitivities of the indicated cells lines to MMS and CPT using the CellTiter-Glo assay. Data are mean ± s.e.m. from 3 biologically independent experiments. **b,** Sensitivities of the indicated hTERT-RPE1 cells lines to MMS and CPT using the CellTiter-Glo assay. Data are mean ± s.e.m. from three biologically independent experiments. **c,** Representative images of fibers from the S1 nuclease experiment. Scale bar: 2 microns.

**Extended Data Fig. 7.**
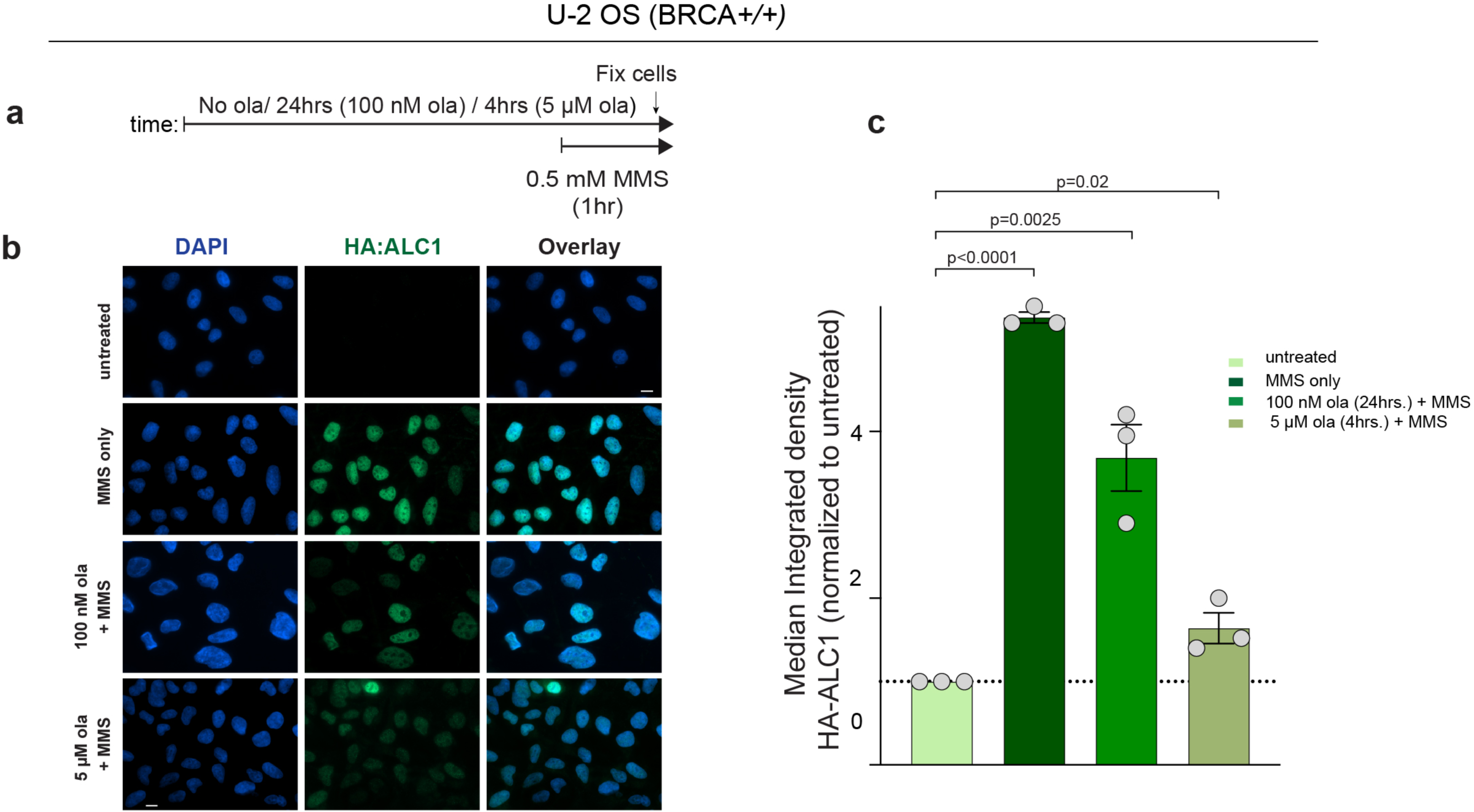
ALC1 is recruited to the damaged chromatin under conditions of reduced PARylation. **a**, Schematic of the experiment. **b-c**, Representative images (**b**) and quantification (**c**) of HA-ALC1 localization to chromatin upon indicated treatments. Scale bar, 10 microns. The median value was normalized to untreated control. Data are mean ± s.e.m. from three biologically independent experiments, *p* value, unpaired Student’s *t*-test. For each experiment, at least 50 cells were analyzed per sample.

**Extended Data Fig. 8.**
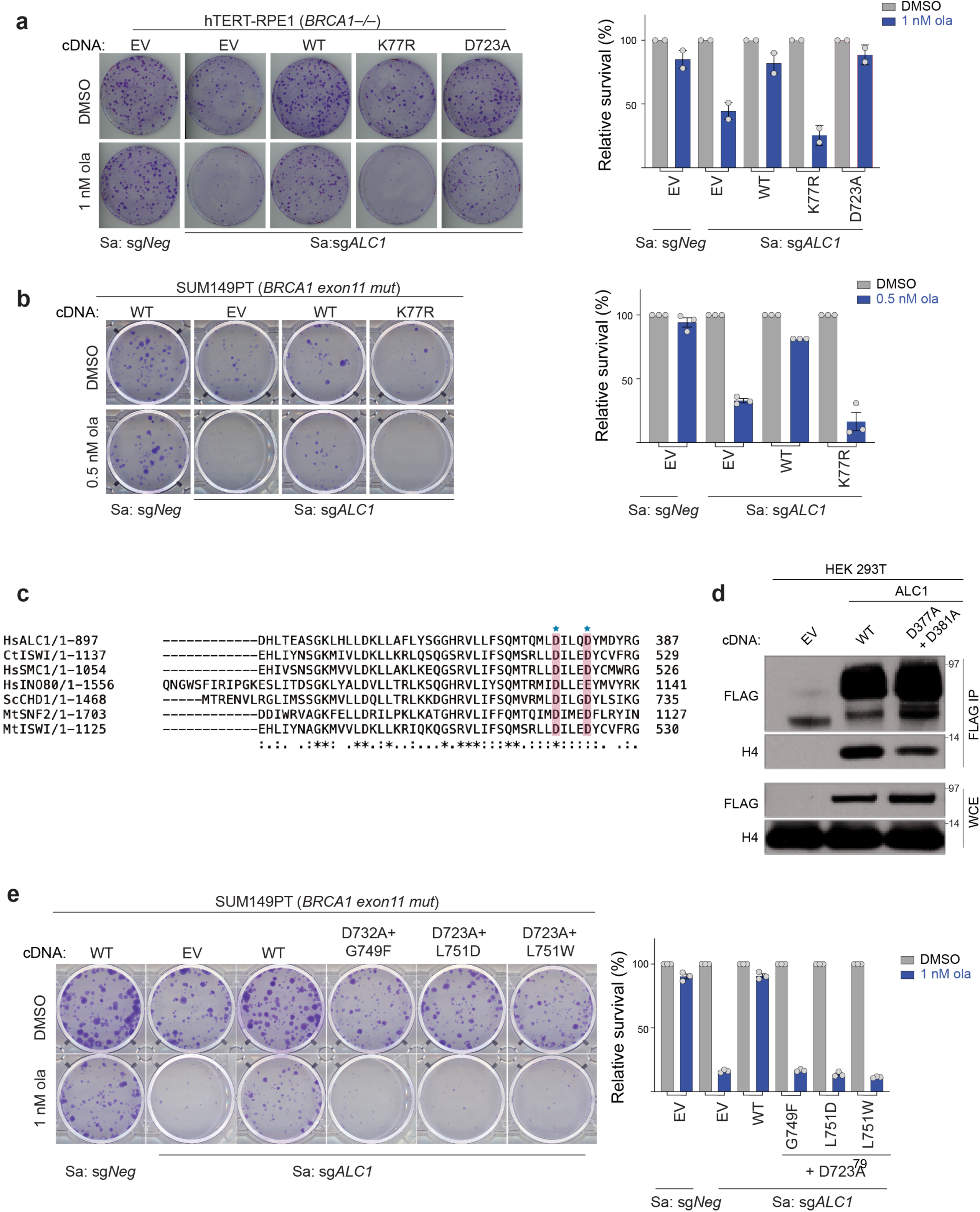
ATPase activity, H4 interaction and macrodomains of ALC1 are essential for protecting BRCA-mutant cells from ola hypersensitivity. **a**, Representative images (left) and quantifications (right) of the clonogenic survival assay using hTERT-RPE1 *BRCA1-/-* cells expressing sg*ALC1* and the indicated ALC1 mutants in ola (1 nM). Data are mean ± s.e.m. from two biologically independent experiments. **b**, Representative images of the clonogenic survival assay (left) and quantification (right) of SUM149PT cells after transduction with sg*ALC1* and ALC1 K77R mutant in ola (0.5 nM). Data are mean ± s.e.m. Number of colonies in the ola treated condition were normalized with its respective untreated counterpart. **c,** Sequence alignment of various chromatin remodelers using Clustal Omega. Histone H4 interacting residues as predicted by PDB:6PWF are highlighted and marked by a blue star. **d,** Immunoblots showing interactions of FLAG ALC1-WT and FLAG ALC1-D377A+ D382A with histone H4. **e**, Representative images of the clonogenic survival assay (left) and quantification (right) of SUM149PT cells expressing sg*ALC1* and indicated ALC1 macro domain mutants in ola (1 nM). Data are mean ± s.e.m. Number of colonies in the ola treated condition were normalized with its respective untreated counterpart.

**Extended Data Fig. 9.**
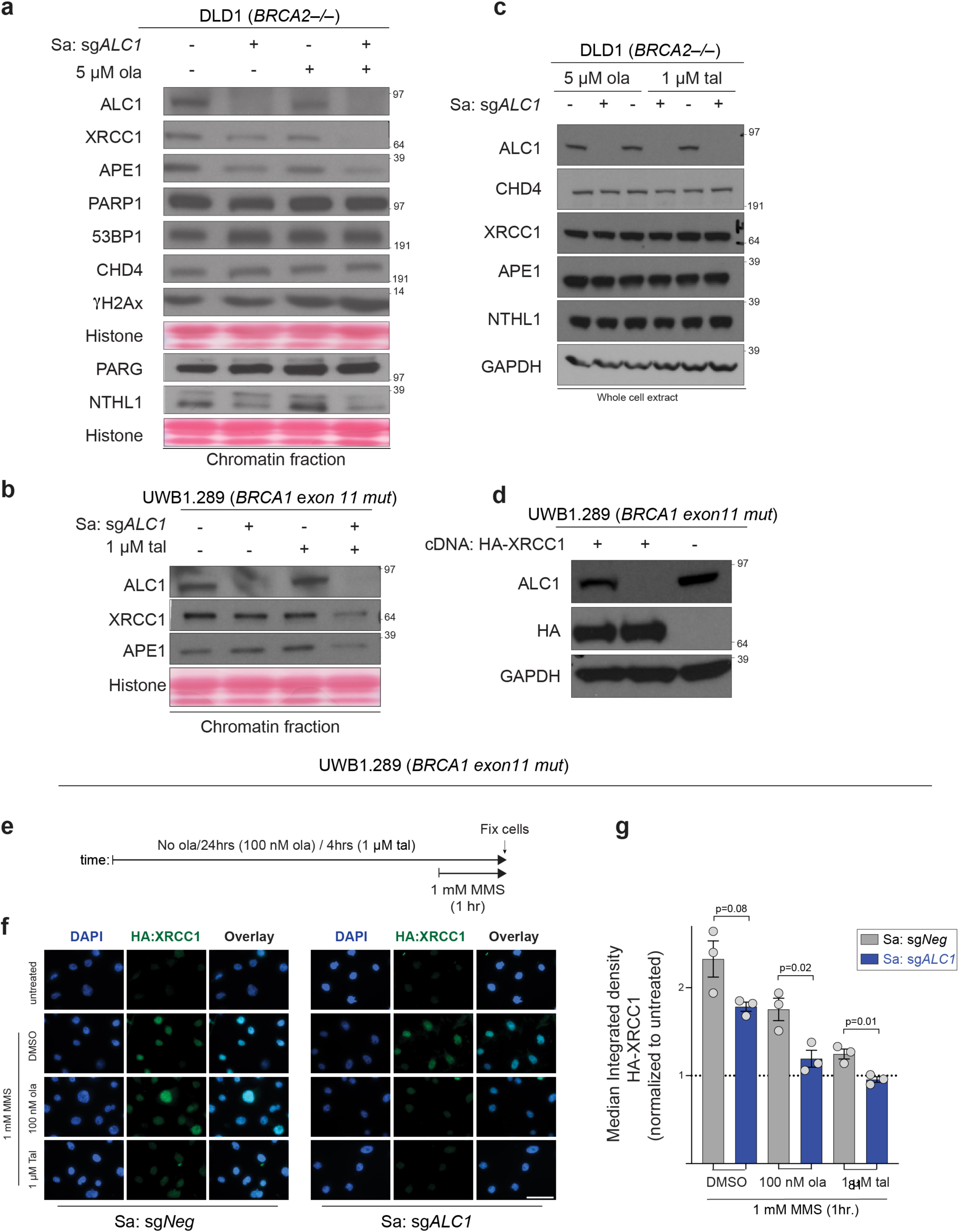
ALC1 co-operates with PARP activity to permit association of repair proteins with chromatin. **a-b**, Cells were fractionated and the chromatin-bound proteins isolated from the indicated DLD1 *BRCA2–/–* **(a)** and UWB1.289 **(b)** cells were immunoblotted. Cells were treated with indicated PARPi for 4 hrs. The data for BER factors is a representative image of at least three biologically independent experiments. Data for *DLD1 BRCA2–/–* cells **(a)** are from the same sample, from two different western blots and the histone levels for each blot are shown by the ponceau staining. **c,** Immunoblot of whole cell lysates of *DLD1 BRCA2-/-* cells showing levels of various proteins upon ALC1 depletion and PARPi treatment. Cells were treated with indicated PARPi for 4 hours. **d,** Immunoblot showing expression levels of HA-XRCC1. **e**, Schematic of the IF experiment. **f-g**, Representative images (**f**) and quantification (**g**) of HA-XRCC1 localization to chromatin upon indicated treatments. Scale bar, 50 microns. Data are mean ± s.e.m. from three biologically independent experiments, *p* value, unpaired Student’s *t*-test. For each cell line, the median value upon MMS treatment was normalized to its respective untreated control.

**Extended Data Fig. 10.**
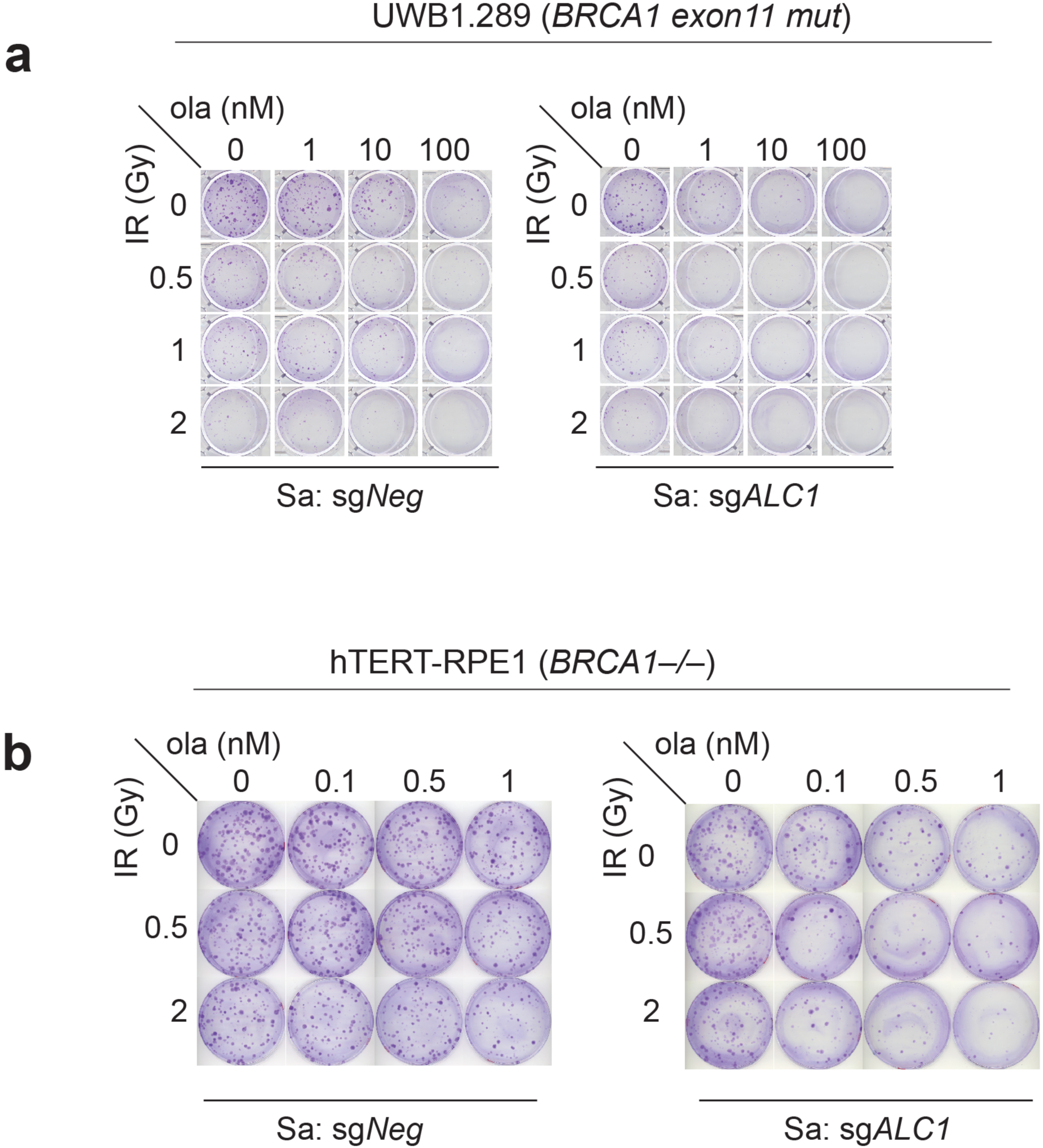
ALC1 loss synergistically enhances IR sensitivity at low olaparib doses. **a,b** Representative images of the clonogenic survival assay to monitor the effect of combining low doses of ola and IR upon ALC1 depletion in UWB1.289 (**a**) and hTERT-RPE1 *BRCA1-/-* cells (**b**).

